# Opposing GIPR brainstem circuits differentially control feeding behaviour

**DOI:** 10.64898/2026.07.01.735388

**Authors:** Natalie S. Figueredo Burgos, Alejandro Lopez-Cruz, Cecilia Skoug, Anna G. Roberts, Kathryn Xie, Iona Davies, Norio Harada, Nobuya Inagaki, Frank Reimann, Fiona M. Gribble, Ben Jones, Daniel I. Brierley, Stefan Trapp, Zachary A. Knight, Alice E. Adriaenssens

**Affiliations:** Centre for Cardiovascular and Metabolic Neuroscience, Department of Neuroscience, Physiology and Pharmacology, University College London, London, United Kingdom; Howard Hughes Medical Institute, University of California, San Francisco, USA; Section of Endocrinology and Investigative Medicine, Imperial College London, London, UK; Department of Endocrinology and Metabolism, School of Medical Sciences, University of Fukui, Fukui, Japan; Department of Diabetes, Endocrinology and Nutrition, Kyoto University, Kyoto, Japan; Institute of Metabolic Science & MRC Metabolic Disease Unit, University of Cambridge, Cambridge, United Kingdom

## Abstract

Central glucose-dependent insulinotropic polypeptide receptor (GIPR) signalling is required for the efficacy of GIP-based obesity therapeutics, yet how distinct subpopulations of GIPR neurons shape appetite remains undefined. Here we show that GIPR neurons in adjacent brainstem nuclei, the area postrema (AP) and nucleus tractus solitarius (NTS), exert opposing control over ingestion. We find GIPR^AP^ neurons dampen post-ingestive satiation, permitting hyperphagia, whereas GIPR^NTS^ neurons are anorectic. In line with this model, we show *Gipr* expression in AP, but not NTS, neurons is necessary for appetite suppression following GIPR antagonism. Additionally, we reveal that GIPR neurons in the AP and NTS occupy distinct gut-brain circuits, and are differentially sensitive to obesity-driven circuit remodelling. These data offer a framework for understanding how current GIPR agonist and antagonist strategies elicit weight loss.

## INTRODUCTION

Glucose-dependent insulinotropic polypeptide (GIP) and Glucagon-like peptide-1 (GLP-1) are gut hormones classically recognized for their role in potentiating postprandial insulin secretion (Holst et al., 1987; Lauritsen et al., 1980). The advent of novel obesity therapies based on both GIP and GLP-1, however, has ignited a keen focus on understanding how the central engagement of their cognate receptors, GIPR and GLP-1R, respectively, affects energy balance (Müller et al., 2025). Whereas the role of central GLP-1R signalling in the control of ingestive behaviour has been extensively characterized, the contribution of GIPR signalling to the regulation of feeding and appetite has perplexed and divided the field (Adriaenssens, 2025; Campbell & Drucker, 2013; Finan et al., 2016; Müller et al., 2018). Notably, both GIPR agonism and antagonism produce clinically meaningful weight loss, suggesting that mechanistically distinct yet equally therapeutically advantageous pathways are engaged by each pharmacological strategy (Frías et al., 2021; Jastreboff et al., 2025).

Core to both therapeutic approaches is the engagement of GIPR neurocircuits in the brain (Gutgesell et al., 2025; Liskiewicz et al., 2023; Liu et al., 2025; Zhang et al., 2021). We and others have previously shown that the dorsal vagal complex (DVC) is a key site of GIPR neuron localisation, activation, and access to peripherally-administered incretin pharmacology (Adriaenssens et al., 2019; Adriaenssens et al., 2023; Borner et al., 2021; de Bray et al., 2025; Liu et al., 2025; Samms et al., 2022). GIPR neurons within the DVC are therefore well placed to integrate peripheral information originating from endocrine and vagal signalling, and make up a critical site of action for the weight-lowering effects of incretin-based pharmacotherapies. Recent transcriptomic analyses demonstrate that GIPR agonism and antagonism elicit distinct differential gene expression profiles in dorsal vagal neurons (Gutgesell et al., 2025), suggesting that GIP-recruited hindbrain circuits may comprise dissociable neural substrates, potentially providing a mechanistic key for resolving the current dichotomy of GIP-based pharmacology. However, the crucial circuit-level understanding of how discrete brainstem GIPR neuronal populations mediate the effects of GIP-based therapies is currently lacking.

To address this knowledge gap, we have defined the anatomical connectivity and functional roles of GIPR neurons in two adjacent brainstem nuclei—the area postrema (AP) and nucleus tractus solitarius (NTS). Using region-specific chemogenetic manipulation and localised *Gipr* deletion models, we show that AP and NTS GIPR neurons exert opposing effects on ingestive behaviour, and that pharmacological GIPR antagonism requires AP *Gipr* expression to supress appetite. We confirm that transcriptomic profiles defining AP or NTS GIPR neurons are distinct, suggesting different molecular mechanisms employed by these cell types. Using a combination of viral circuit mapping and *in vivo* neuronal activity recordings, we show that GIPR AP and NTS neurons are engaged by largely distinct vagal afferent populations that integrate discrete interoceptive signals from the gut. These circuits are differentially disrupted in response to Western diet-induced obesity, providing important context for their relative pathophysiological relevance. Collectively our data reveal that GIPR^AP^ and GIPR^NTS^ neurons represent opposing circuits, and that defining the balance of how these populations are engaged is essential to building a coherent model of how current and future GIPR agonists or antagonists decrease body weight.

## RESULTS

### GIPR Neurons in the Area Postrema and Nucleus Tractus Solitarius Oppositely Regulate Ingestion

To date, studies aiming to characterise the role of AP and NTS GIPR neurons in appetite have either applied stereotaxic manipulations to the entire DVC (Adriaenssens et al., 2023), or have generated loss of function models using pan-neuronal Cre drivers. Transcriptomic analysis of murine DVC snRNAseq data, however, revealed that GIPR neurons in the AP and the NTS exhibit unique molecular profiles (Fig 1A-B). While both AP and NTS GIPR neurons expressed markers for inhibitory cell types, GIPR neurons in the AP (GIPR^AP^) expressed higher levels of markers associated with GABAergic neurons (Fig 1B). GIPR neurons in the NTS (GIPR^NTS^) exhibited a mixed profile, demonstrating high expression levels of glycinergic neuronal markers as well as markers for excitatory cell types (Fig 1A-B). These Data indicate that GIPR^AP^ and GIPR^NTS^ neurons may utilise different signalling mechanisms and recruit distinct circuits to control energy homeostasis.

**Figure 1:**
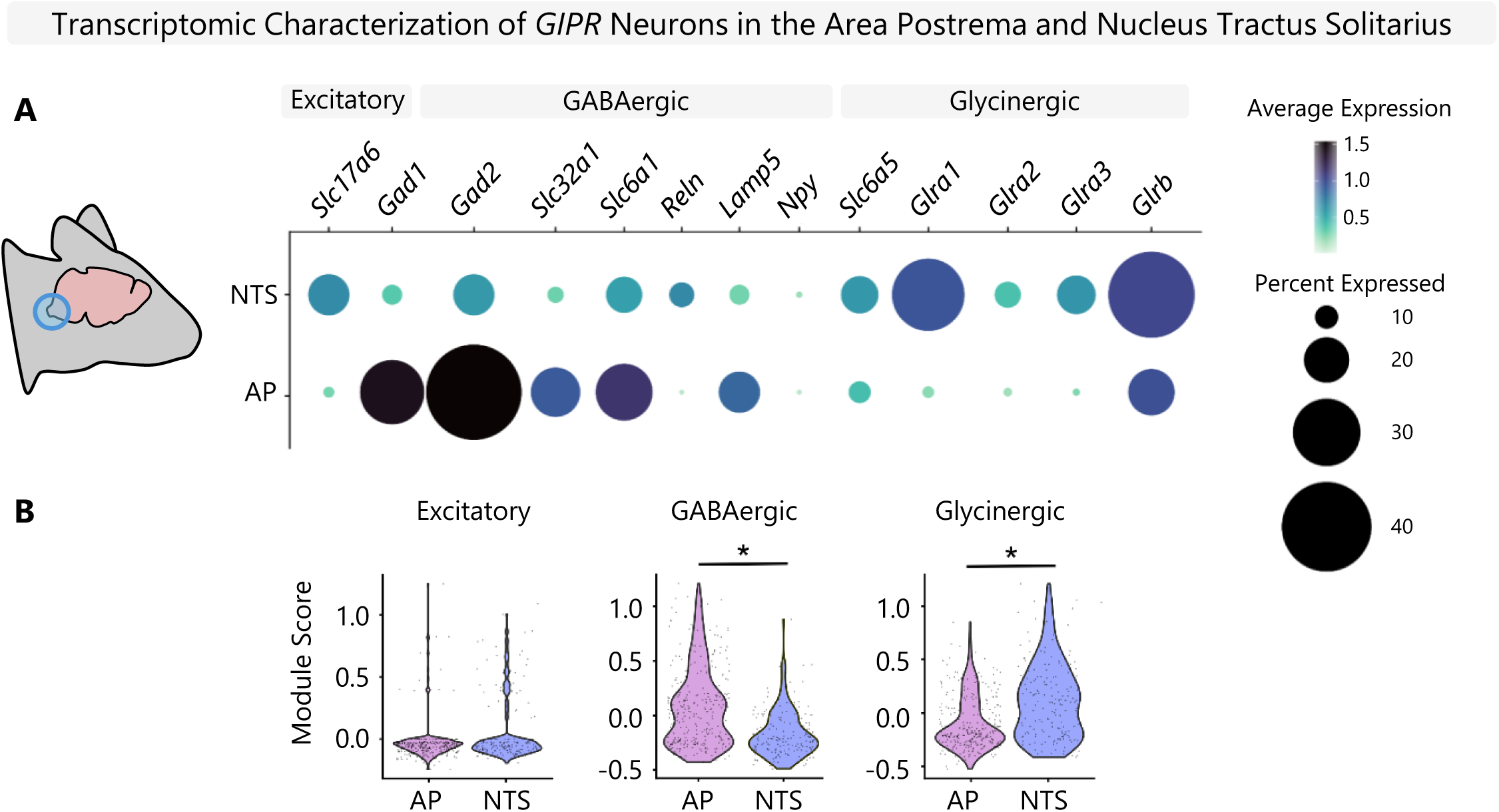
Transcriptomic Characterization of GIPR Brainstem Neurons in Mice. (A-B) Transcriptomic analysis of *Gipr+* neurons in the dorsal vagal complex was conducted using publicly available snRNAseq datasets generated from DVC tissue isolated from wildtype mice. (A) Dotplot of neuron class markers expressed across murine AP and NTS *Gipr* neurons. (B) Module scores representing relative enrichment of GABAergic, glutamatergic (excitatory) and glycinergic neuronal markers in GIPR^AP^ versus GIPR^NTS^ neurons isolated from mouse DVC.

Given these differences in molecular makeup between GIPR AP and NTS neurons, approaches that fail to target each population independently may miss the relative contribution of GIPR^AP^ versus GIPR^NTS^ cells to the regulation of ingestion. We therefore optimized neuronal targeting protocols to allow us to define the functional roles of GIPR^AP^ versus GIPR^NTS^ neurons in the regulation of food intake.

We selectively expressed a Cre-inducible Gq-coupled designer receptor exclusively activated by designer drugs (DREADD), hM3Dq (Alexander et al., 2009), in either the AP (Fig 2A) or NTS (Fig 2D) of *Gipr*-Cre mice (Adriaenssens et al., 2019). Food intake was assessed using a counterbalanced crossover design with FED3 monitoring in the home cages (Matikainen-Ankney et al., 2021). Acute activation of GIPR^AP^ neurons unexpectedly revealed that this population comprises a previously uncharacterized feeding-promoting GIPR circuit, driving a transient increase in food intake during the first 4 hours of the dark phase (Fig 2B), consistent with the half-life of CNO. Meal pattern analysis showed that this orexigenic effect correlated with increased meal size and meal duration (Fig 2C), consistent with reduced satiation (Rathod & Di Fulvio, 2021; Zorrilla et al., 2005). In contrast, activation of GIPR^NTS^ neurons produced a sustained suppression of food intake (Fig 2E) associated with reduced meal number (Fig 2F), indicating enhanced satiety (Rathod & Di Fulvio, 2021; Zorrilla et al., 2005).

**Figure 2:**
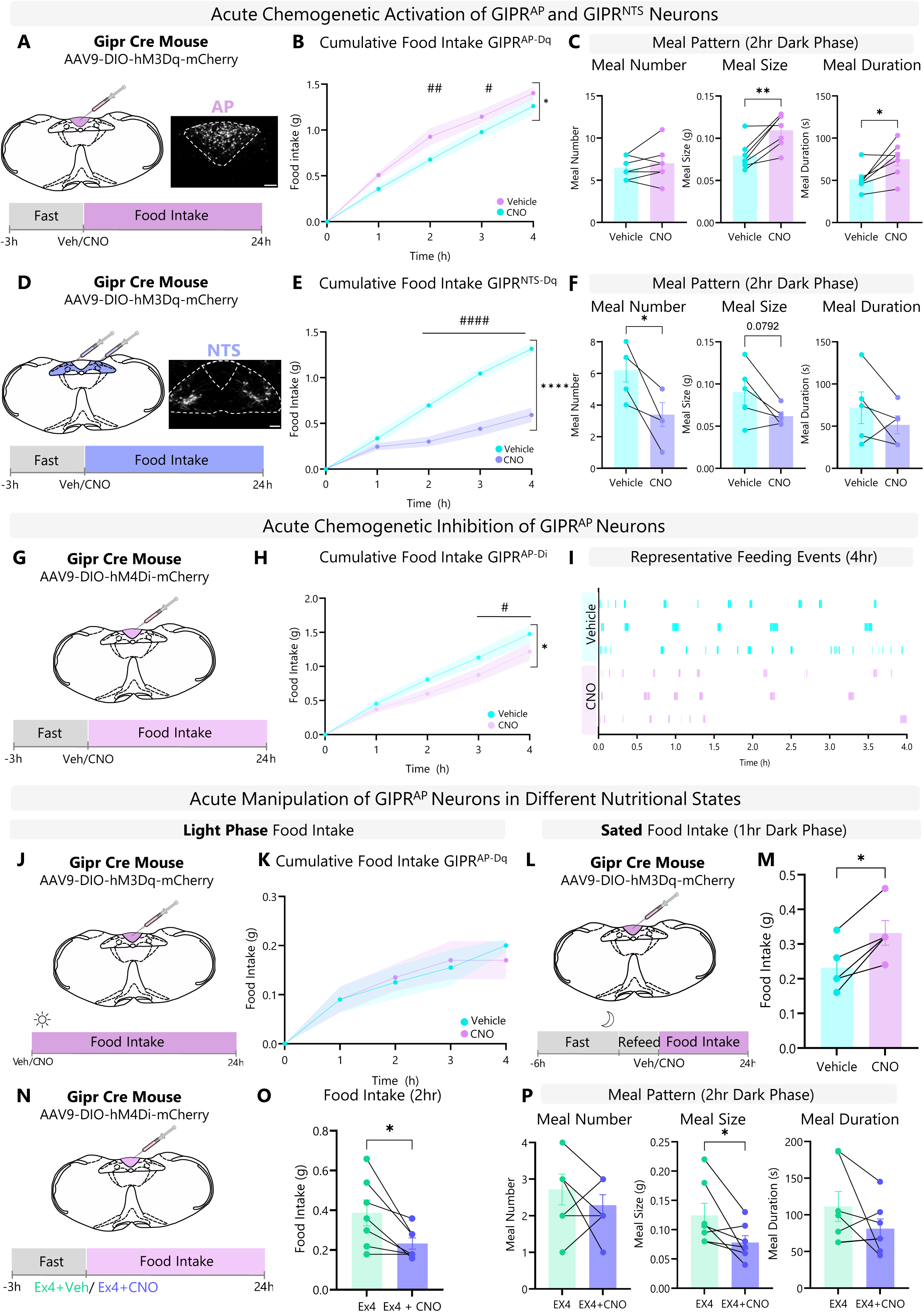
GIPR ^AP^ and GIPR ^NTS^ Neurons Oppositely Regulate Food Intake. A-F, Gipr-Cre mice were injected with AAV-hSyn-DIO-hM3D(Gq)-mCherry into the AP (A) or NTS (D) to produce GIPR^AP-Dq^, or GIPR^NTS-Dq^ mice, respectively. G-I, Gipr-Cre mice were injected with AAV-hSyn-DIO-hM4D(Gi)-mCherry into the AP to produce GIPR^AP-Di^ mice. Mice were housed in home cages equipped with FED3 pellet dispensers. CNO (1 mg/kg) or vehicle was injected intraperitoneally (i.p.) at the onset of the dark phase and changes to feeding behaviour were measured (B-C, E-F, H-I). J-M, For nutritional state experiments, GIPR^AP-Dq^ mice received CNO (1 mg/kg) or vehicle (i.p.) in the light phase (ZT7) (J) or after a 6hr fast followed by 1hr ad lib access to food for sated paradigm (L). N-P, Ex4 + Veh or Ex4 + CNO were administered i.p. to GIPR^AP-Di^ mice at onset of dark phase to assess effect of inhibition of GIPR^AP^ on EX4-induced anorexia. Data are plotted as means +/-SEM. Statistical comparisons made using a repeated measures 2-way ANOVA with a Sidak’s post-hoc test, paired or unpaired t-test as appropriate are indicated by * symbols. # symbols indicate significance from multiple comparisons of vehicle vs. CNO over hourly food intake. # p < 0.05, ## p < 0.01, #### p < 0.001, * p < 0.05, ** p < 0.01, **** p < 0.001; n = 4-7, Scale bar: 100 µm

Given our unexpected finding that GIPR^AP^ neurons increase food intake, we sought to determine whether activity of GIPR^AP^ neurons is required to maintain feeding drive during normal meal intake periods. We acutely silenced this population at the onset of the dark phase using hm4Di DREADDs (Fig 2 G-I). Inhibiting GIPR^AP^ neurons at the onset of feeding resulted in significant reduction in dark phase food intake (Fig 2H-I), suggesting that GIPR^AP^ neurons support ingestive drive in the dark phase.

Together, these data indicate that while populations of neurons in both the AP and the NTS are equipped to respond to GIP, their activity exerts opposing effects on feeding through distinct behavioural mechanisms.

### GIPR Area Postrema Neurons Promote Feeding by Dampening Post-Ingestive Satiation Signals

To further define the hyperphagic drive of GIPR^AP^ neurons, we examined the effects of acute manipulation of this population across different nutritional states. We first performed chemogenetics-assisted phenotyping in the light phase (Fig 2J) during which food intake is typically low in mice (Ellacott et al., 2010). Under these conditions, activation of GIPR^AP^ neurons did not significantly alter intake, indicating that recruitment of GIPR^AP^neurons does not evoke sufficient drive to eat outside of regular feeding periods (Fig 2K).

We next used a post-satiation paradigm specifically designed to interrogate changes to orexigenic drive in full animals (Fig 2L-M). Following a 6 h fast that extended 1 h into the dark phase, mice were given ad libitum access to food for 1 h, after which mice were dosed with either vehicle or CNO, and subsequent feeding behaviour was assessed in the sated state. No difference was observed in the amount of food eaten during the 1 h refeed prior to vehicle/CNO administration (Supp Fig 1). Post-refeed, acute activation of GIPR^AP^ neurons significantly increased food intake during the first hour after stimulation (Fig 2M).

Data from our post-satiation paradigm suggest that inhibition of GIPR^AP^ neurons should enhance the anorectic effect of known post-ingestive termination signals. To test this, we inhibited GIPR^AP^ neurons while pharmacologically activating GLP1R circuits using Exendin-4 (Ex4) (Fig 2N-P, Supp Fig 2). Combined GIPR^AP^ inhibition and Ex4 treatment produced a greater reduction in 2-hour food intake (Fig 2O), and enhanced Ex4-mediated suppression of meal size compared to Ex4 alone (Fig 2P). These findings indicate that GIPR^AP^ neurons interact with satiation-promoting GLP-1R circuits.

Collectively these data suggest that GIPR^AP^ circuits promote feeding by dampening post-ingestive satiation signals and suppressing downstream meal termination pathways.

### *Gipr* Expression in the AP and NTS is Necessary for Maintaining Normal Feeding Patterns

Next, we interrogated the necessity of *Gipr* expression in either the AP or the NTS for regulating feeding behaviour. We generated region-specific *Gipr* deletion models by delivering an AAV encoding Cre recombinase into either the AP or NTS of *Gipr*^fl/fl^ mice (Joo et al., 2017), producing *Gipr*^AP-KO^ or *Gipr*^NTS-KO^ animals, respectively. Sham control animals were injected with an AAV-mCherry vector. RNAscope quantification of *Gipr* mRNA confirmed 51.7% and 47% knock down in *Gipr*^AP-KO^ (Fig 3A-B) and *Gipr*^NTS-KO^ (Fig 3D-F) animals, respectively. In alignment with our chemogenetic models, deletion of *Gipr* in the AP reduced meal duration under sated conditions, indicating that selective loss of GIPR signalling in the AP enhances satiation (Fig 3C). *Gipr* deletion in the NTS increased dark phase meal duration, indicating reduced satiation (Fig 3F). These reciprocal effects produced by region-specific *Gipr* deletion indicate that endogenous GIP signalling engages these parallel hindbrain circuits to differentially affect meal structure. These data further support a role for anatomically distinct GIPR neuronal populations within the dorsal vagal complex that exert opposing effects on feeding: GIPR^AP^ neurons increase food intake by reducing satiation, whereas GIPR^NTS^ neurons suppress intake through enhanced satiety. In accordance with this model, *Gipr*^AP-KO^ mice maintained on a chow diet trended towards decreased 21-hr food intake compared to sham controls, while *Gipr*^NTS-KO^ animals trended toward net increase in food intake (Supp Fig 3).

**Figure 3:**
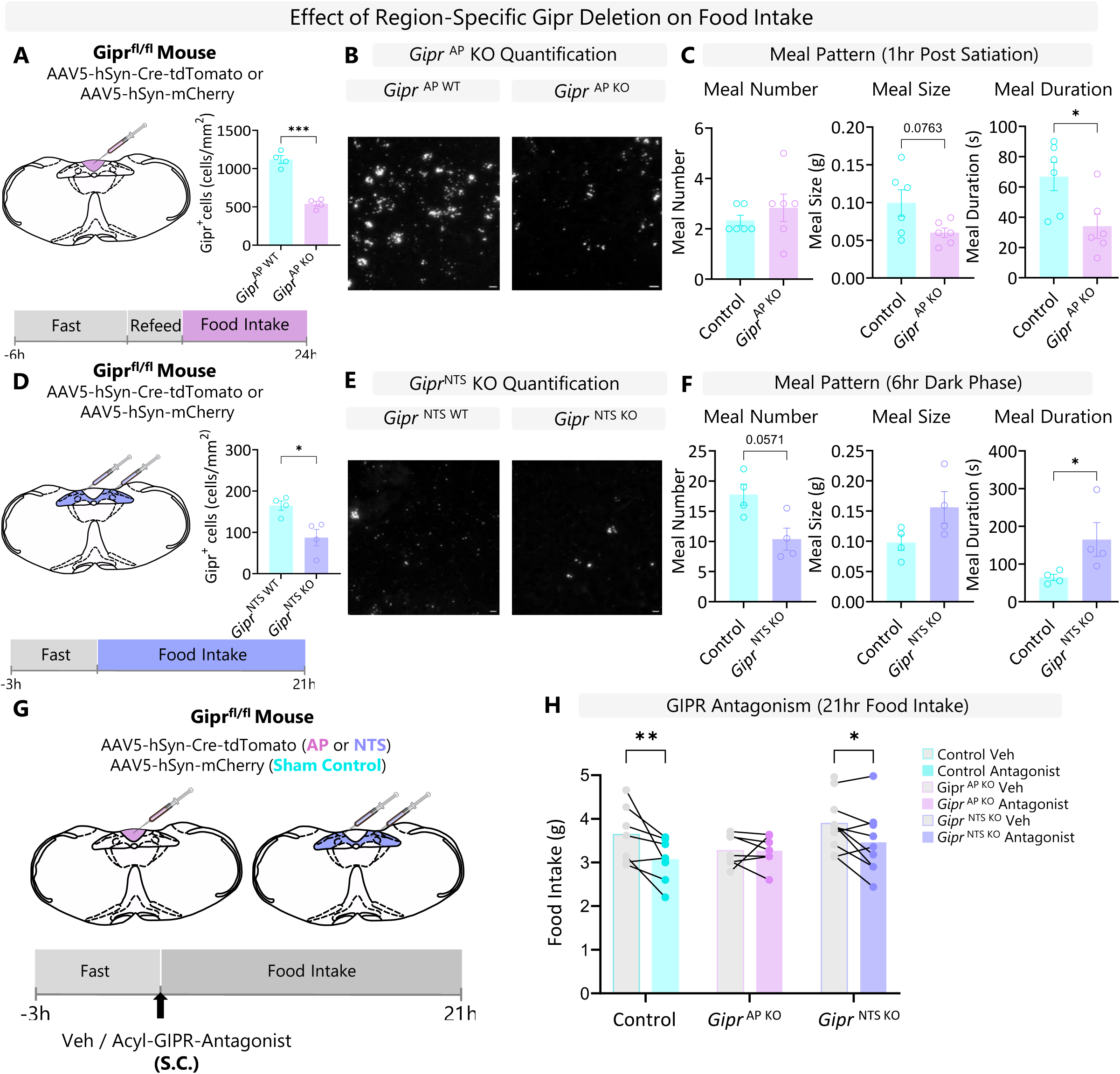
G*i*pr Expression in the AP and NTS is Necessary for Maintaining Normal Feeding Patterns and *Gipr* Expression in the AP is Necessary for Anorectic Effects of Pharmacological GIPR Antagonism. A-H, Gipr^fl/fl^ mice were injected with AAV-hSyn-Cre-tdTomato to produce selective KO of *Gipr* in the AP (Gipr^AP-KO^) or NTS (Gipr^NTS-KO^). Sham injected mice received a control virus in the AP and NTS. (A-B, D-E) Fluorescent *in situ* hybridization against *Gipr* was used to quantify degree of knockout. Baseline changes in food intake in sated paradigm (C) or dark phase feeding (F) were measured using FED3 devices in home cages. C, For sated paradigm, Gipr^AP-KO^ and control mice were fasted for 6hr and given ad libitum access to food for an hour, changes in food intake patters were then assessed. F, For dark phase paradigm Gipr^NTS-KO^ and control mice were fasted for 3h of the light phase and FED3 devices were returned at onset of dark phase. G-H, To assess necessity of *Gipr* in the AP or NTS for the therapeutic effect of pharmacological GIPR antagonism, mice were administered with Acyl-GIPR-Antagonist (3µmol/kg) subcutaneously 15 minutes prior to dark onset. Changes in food intake were assessed for 21hr using FED3 devices (H). Data are plotted as means +/-SEM. Statistical comparisons made using a repeated measures 2-way ANOVA with a Sidak’s post-hoc test, paired or unpaired t-test as appropriate. * p < 0.05, ** p < 0.01, **** p < 0.001; n = 4-7, Scale bar: 10 µm

### Selective loss of *Gipr* Expression in AP Neurons Attenuates Appetite Suppression in Response to Pharmacological GIPR Antagonism

Given the unique orexigenic influence exerted by GIPR^AP^ neurons compared to other GIPR circuits, we hypothesized that GIPR neurons in the AP could have an important role in mediating the appetite-suppressing effects of pharmacological GIPR antagonists. To test this hypothesis, we administered acylated peptide GIPR antagonist (Yang et al., 2022) into sham control, GIPR^AP-KO^ and *Gipr*^NTS-KO^ animals at the onset of the dark phase (Fig 3I). GIPR antagonism elicited robust suppression of 21 h chow intake in sham control and *Gipr*^NTS-KO^ animals. (Fig 2J) However, GIPR antagonist-induced appetite suppression was ablated in GIPR^AP-KO^ animals (Fig 3J). These data identify GIPR^AP-KO^ neurons as critical mediators of the weight lowering effects of GIPR antagonists.

### GIPR Neurons in the Area Postrema and Nucleus Tractus Solitarius Receive Distinct Regulatory Input

Given the contrasting roles GIPR^AP^ and GIPR^NTS^ neurons in controlling feeding behaviour, we sought to define the presynaptic partners that differentially engage these circuits. To map the central and peripheral inputs regulating GIPR^AP^ and GIPR^NTS^ neuron activity, we performed rabies-assisted monosynaptic input tracing. Both GIPR^AP^ and GIPR^NTS^ neurons received descending inputs from the bed nucleus of the stria terminalis (BNST), the paraventricular nucleus (PVH) and the central amygdala (CeA), although with distinct relative densities. GIPR^AP^ neurons additionally received inputs from the retrochiasmatic nucleus (Rch), periaqueductal grey (PAG), pontine nucleus (Pn) and zona incerta (Zi), whereas GIPR^NTS^ neurons were preferentially innervated by the area postrema and parasubthalamic nucleus (pSTH) (Fig 4).

**Figure 4:**
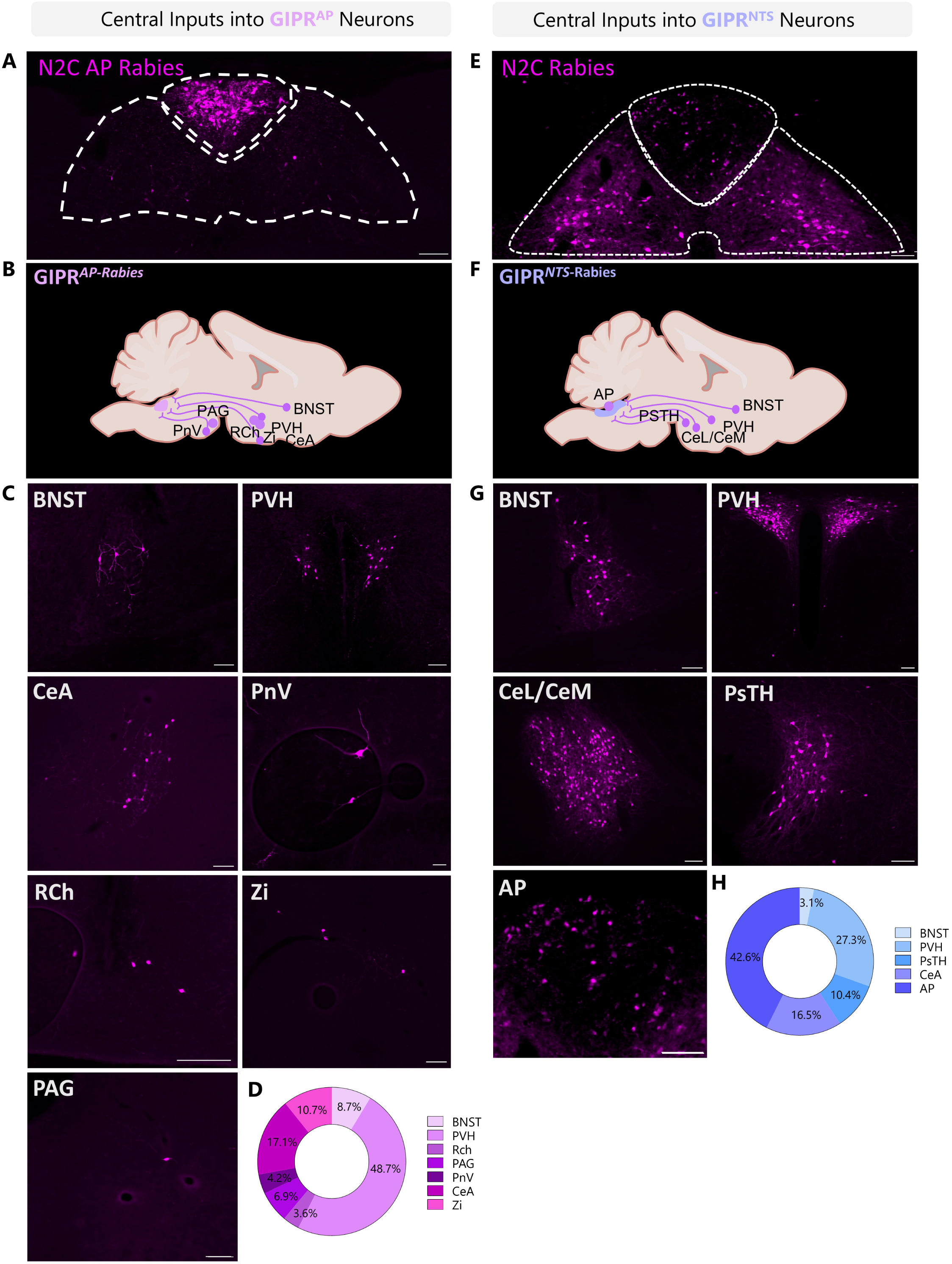
Central Monosynaptic Input Tracing to GIPR ^AP^ vs GIPR ^NTS^ Neurons. Gipr-Cre mice were injected with AAV-TREtight-mTagBFP2-N2cG and AAV-hSyn-FLEX-splitTVA-EGFP-Tta helper virus into the AP or NTS. Subsequently, mice were injected with EnvA-N2C-mCherry rabies. Central regulatory inputs were mapped. (A-C) GIPR^AP^ neurons receive central innervation from the bed nucleus of the stria terminalis (BSNT), paraventricular nucleus hypothalamus (PVH), pontine nucleus (PnV), retrochiasmatic nucelus (RcH), zona incerta (ZI), central amygdala (CeA), and periaqueductal grey (PAG), quantified in (D). (E-G) GIPR^NTS^ neurons receive central monosynaptic inputs from the area postrema (AP), BNST, parasubthalamic nucleus (psTH), CeA and PVH quantified in (H). n = 2-4, Scale bar: 100 µm

### Molecular Identity of Vagal Afferents Defines Differential Interoceptive Recruitment of GIPR Neurons

The dorsal vagal complex represents a critical integration hub between the brain and the periphery, with both vagal and spinal circuits conveying vital interoceptive signals into the brainstem to maintain homeostasis (Browning et al., 2017; de Lartigue et al., 2026; Jänig, 1996). As such, we examined the peripheral sensory input to GIPR neurons in the AP and NTS. Both GIPR^AP^ and GIPR^NTS^ neurons received dense innervation from vagal afferent neurons (VANs) (Fig 5A-D). To determine whether these rabies-labelled afferents comprised common or distinct populations, we performed two-colour rabies tracing analysis to compare input overlap. This analysis revealed minimal convergence (Supp Fig 4A), indicating that AP and NTS GIPR neurons are embedded within largely non-overlapping vagal afferent → DVC circuits.

**Figure 5:**
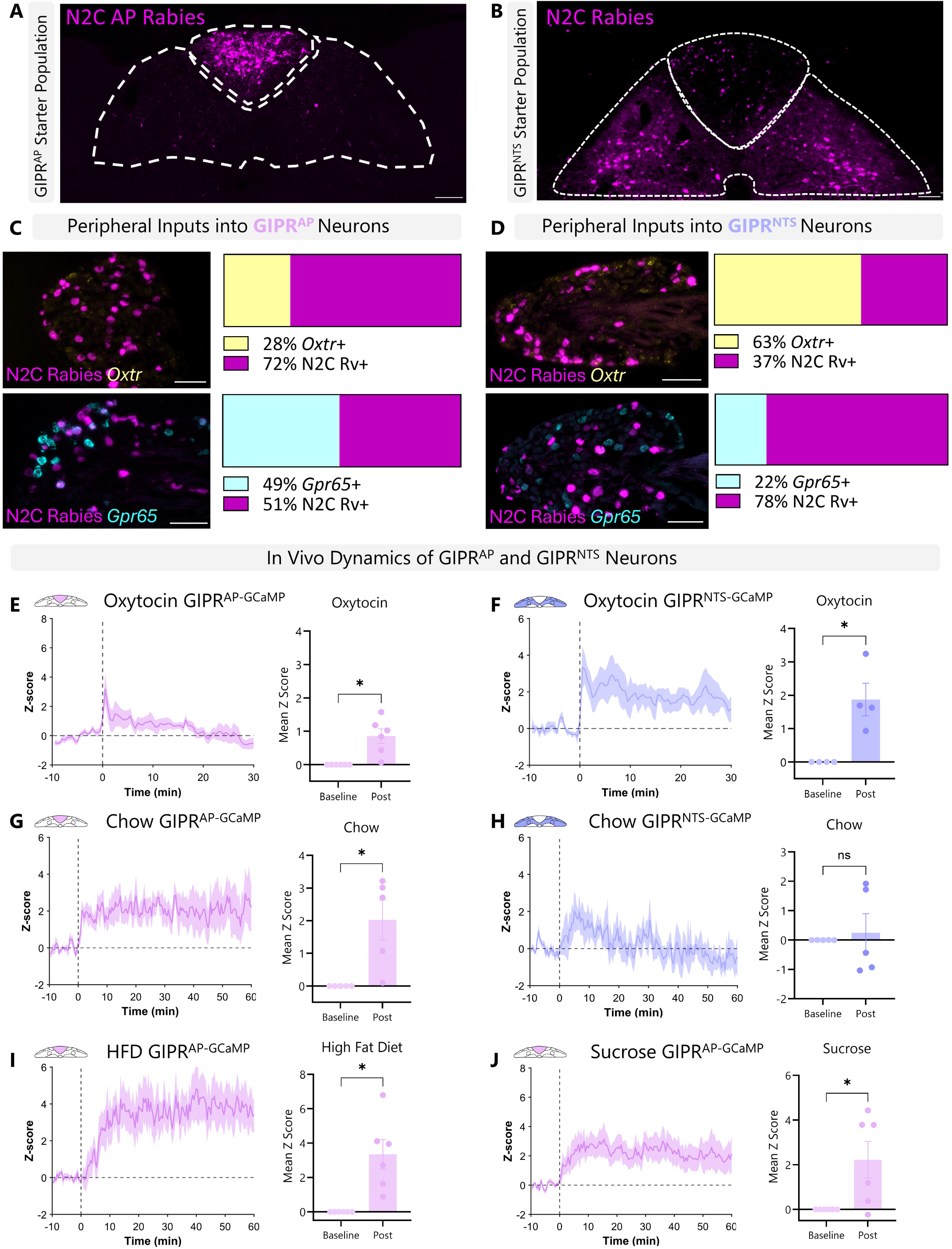
Vagal Inputs Gipr ^AP^ vs Gipr ^NTS^ Define Interoceptive Recruitment of GIPR Neurons *in vivo*. A-B, Gipr-Cre mice were injected with AAV-TREtight-mTagBFP2-N2cG and AAV-hSyn-FLEX-splitTVA-EGFP-Tta helper virus into the AP or NTS. Two weeks later, mice were injected with EnvA-N2C-mCherry rabies in the AP (A) or NTS (B). C-D, Monosynaptic inputs were identified in the nodose ganglion. Molecular characterization of inputs was performed using fluorescent *in situ* hybridization probing for *Oxtr* (yellow) and *Gpr65* (cyan), markers of mechanosensing and chemosensing vagal afferents, respectively. (C) Gipr ^AP^ neurons receive inputs from vagal afferents of which 49% are *Gpr65*+ and 28% are *Oxtr*+. (D) Vagal inputs to Gipr ^NTS^ neurons are 63% *Oxtr*+ and 22% *Gpr65*+. Gipr-Cre mice were injected with AAV-CAG-Flex-GCaMP6s and implanted with photometry fiber targeting either the AP or NTS. GIPR^AP^ (E) and GIPR^NTS^ (F) responses to IP injection of oxytocin. GIPR^AP^ (G) and GIPR^NTS^ (H) responses to self-paced chow consumption. GIPR^AP^ response to high fat diet (I) and sucrose (J). Data are plotted as means +/-SEM. Statistical comparisons made using paired t-test as appropriate. * p < 0.05; n = 4-6, Scale bar: 100 µm

To define the interoceptive signals conveyed by labelled vagal inputs to GIPR neurons, we performed RNAscope using molecular identifiers for mechanosensory and chemosensory vagal afferent populations. *Oxtr* and *Glp1r* are established markers for predominantly mechanosensory VAN populations, whereas *Gpr65* expression denotes chemosensory VANs (Bai et al., 2019; Williams et al., 2016). Only 11% of vagal inputs to GIPR^DVC^ cells expressed *Glp1r* (Supp Fig 4B), so we decided to take *Oxtr* forward as our principal mechanosensory VAN marker. We confirmed that *Piezo2* expression is enriched in *Oxtr*+ versus *Gpr65*+ VANs (Supp Fig 4C), consistent with a mechanosensory cell type. 49% of traced vagal neurons innervating GIPR^AP^ neurons expressed *Gpr65*, whereas 28% expressed *Oxtr* (Fig 5C). In contrast, inputs to GIPR^NTS^ neurons were enriched for *Oxtr*-expressing vagal afferents (63%), with only 22% expressing *Gpr65* (Fig 5D). These data indicate that mechanosensory vagal afferents predominantly engage GIPR^NTS^ neurons, whereas GIPR^AP^ neurons receive a relatively greater proportion of chemosensory vagal input. Collectively, these findings suggest that distinct vagal afferent populations encoding different interoceptive modalities recruit anatomically and functionally segregated GIPR neuron populations in the AP and NTS.

To test whether this circuit organization is reflected *in vivo*, we performed fiber photometry in freely-behaving animals to monitor calcium dynamics in either GIPR^AP^ or GIPR^NTS^ neurons in response to defined physiological stimuli. We first administered exogenous oxytocin to engage OXTR associated vagal signalling. Consistent with the presence of *Oxtr*+ inputs to both GIPR^AP^ and GIPR^NTS^ neurons, oxytocin induced significant increase in calcium responses in AP (Fig 5E) and NTS (Fig 5F) populations. We next assessed responses to nutritional stimuli, key chemosensory signals that regulate VANs (Bai et al., 2019; Williams et al., 2016). Ingestion of standard chow evoked a sustained increase in activity in GIPR^AP^ neurons (Fig 5G), but failed to elicit significant activation of GIPR^NTS^ neurons (Fig 5H). We observed nutrient-simulated activation in GIPR^AP^ neurons when mice were challenged with both a high-calorie mixed diet (60% HFD) (Fig 5I) or liquid sucrose (Fig 5J).

Together, these findings demonstrate that GIPR^AP^ and GIPR^NTS^ neurons exhibit distinct response profiles *in vivo* to interoceptive stimuli. Specifically, GIPR^AP^ neurons respond to dietary and chemosensory cues, whereas GIPR^NTS^ neurons preferentially respond to oxytocin-engaged vagal signalling. Collectively, these data establish that modality-specific vagal afferent organization is functionally reflected in the differential recruitment of brainstem GIPR neuron populations in vivo.

### Western Diet Remodels Inputs to GIPR Neurons

Recent studies have shown that the networking and connectivity of appetite regulatory neurons are highly plastic in response to metabolic stressors (Beutler et al., 2020, Linehan et al., 2020, Ferrario et al., 2024). This maladaptive plasticity blunts interoception of post-ingestive signals and nutritional state (Kentish et al., 2012; Page & Kentish, 2017). We thus sought to determine whether GIPR neurocircuits in the brainstem undergo similar maladaptive rewiring in response to obesogenic conditions.

Aged-matched mice were maintained on either a Western diet (WD) or control chow diet for 10 weeks. Body composition analysis and glucose tolerance testing demonstrated significant metabolic dysfunction in mice fed WD, with increased fat mass and impaired glucose tolerance compared to the control-fed group (Supp Fig 5). We next performed rabies-assisted monosynaptic input tracing to determine whether central and peripheral inputs to GIPR^AP^ and GIPR^NTS^ neurons change in response to WD (Fig 6). While vagal innervation of GIPR^AP^ neurons was largely preserved (Fig 6A-B,E-F), Western diet exposure produced a robust reduction in vagal afferent inputs to GIPR^NTS^ neurons (Fig 6C-D, G-H). Deafferentation of GIPR^NTS^ neurons was driven by selective loss of *Oxtr*+ mechanosensory VANs (Fig 6H). These findings indicate that GIPR^NTS^ neurons undergo selective loss of mechanosensory vagal connectivity in an obesogenic environment, potentially reducing recruitment by peripheral interoceptive signals. In contrast, peripheral inputs to GIPR^AP^ neurons appear resistant to obesity-associated circuit remodelling.

**Figure 6:**
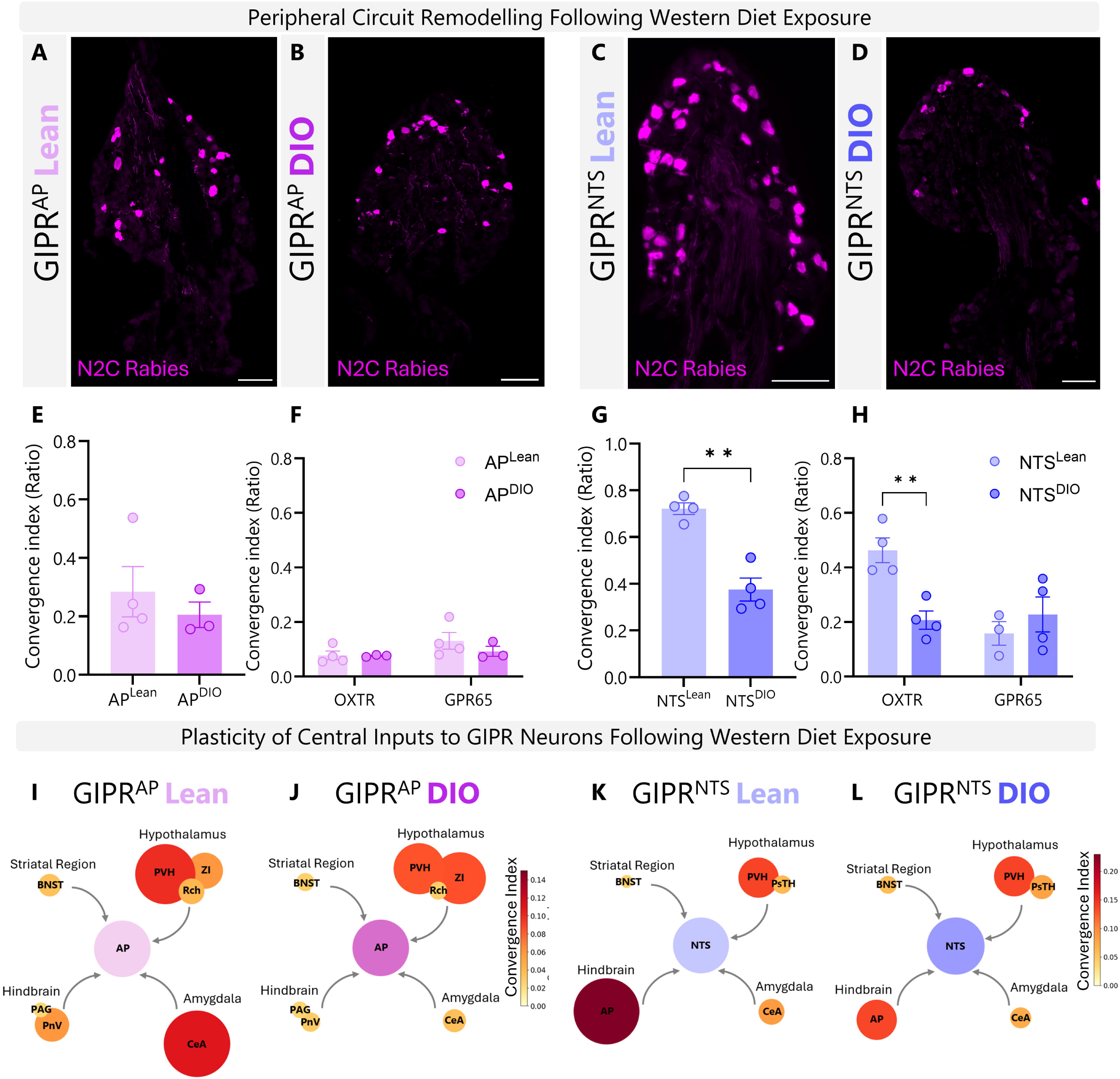
**GIPR Circuit Remodelling Following Exposure to Western Diet**. Gipr-Cre mice were fed for 8-10 weeks with control diet or Western diet before being injected with AAV-TREtight-mTagBFP2-N2cG and AAV-hSyn-FLEX-splitTVA-EGFP-Tta helper virus into the AP or NTS. Two weeks later, mice were injected with EnvA-N2C-mCherry rabies into the AP (A-B) or NTS (C-D). Changes in vagal inputs to GIPR^AP^ or GIPR^NTS^ circuitry were characterised. Convergence index changes between lean and obese vagal inputs were calculated for total vagal inputs and molecularly defined populations for either GIPR^AP^ (E-F) or GIPR^NTS^ (G-H) neurons. Changes to central inputs to GIPR^AP^ (I-J) or GIPR^NTS^ (K-L) neurons in lean vs obese state are represented as heat map dot-plots representing convergence index ratio. Data are plotted as means +/-SEM. n = 3-4, Scale bar: 100 µm

We next assessed changes to central inputs to GIPR^AP^ and GIPR^NTS^ neurons. Western diet exposure redistributed the relative input architecture to GIPR^NTS^ neurons, decreasing AP innervation while increasing the contributions from BNST and pSTH pathways (Fig 6K-L). Similarly, the relative distribution of central inputs to GIPR^AP^ remodelled when comparing lean and obese states (Fig 6 I-J). The proportion of inputs from PVH, CeA, Rch, BNST and PnV decreased whilst from Zi and PAG increased.

Collectively, these findings evidence maladaptive changes to brainstem GIPR neurocircuits as a consequence of western diet exposure.

## DISCUSSION

In this study, we investigated how *Gipr*-expressing neurons located in different brainstem nuclei contribute to the regulation of feeding behaviour. Our findings reveal that GIPR neurons in the AP and NTS are part of distinct anatomical and functional circuits that exert opposing effects on food intake and integrate different interoceptive signals produced during ingestion. We further show that intact sensing of endogenous GIP by both AP and NTS GIPR neurons is necessary for maintaining normal feeding microstructure, and that intact GIPR signalling in AP neurons is necessary for the appetite suppressing effects of pharmacological GIPR antagonism. Lastly, we show that GIPR neurocircuits are susceptible to maladaptive changes in response to obesogenic pressures.

Together, these findings provide a refined framework for understanding how central GIPR signalling in the brainstem contributes to energy balance and informs our understanding of the neuron populations that underpin the effects of different GIP-based approaches for obesity therapeutics.

### GIPR Neurons in the AP and NTS Engage Different Mechanisms to Control Feeding Behaviour

Transcriptomic profiling of DVC GIPR cells in mouse reveals distinct molecular heterogeneity when comparing GIPR^AP^ and GIPR^NTS^ neurons. Here, we demonstrate that this molecular heterogeneity extrapolates to functionally opposing roles for GIPR^AP^ and GIPR^NTS^ cells in the context of ingestive behaviour: GIPR^AP^ neurons exert an orexigenic tone to suppress post-ingestive satiation signals, whereas GIPR^NTS^ neurons enhance satiety.

The orexigenic phenotype of GIPR^AP^ neurons is, to our knowledge, the first description of a GIPR-defined population that promotes feeding. It is, however, mechanistically consistent with the established role of these cells in gating nausea and malaise. We and others have shown that GIPR^AP^ cells are inhibitory and predominantly form dense local arborizations in the AP and neighbouring NTS (Adriaenssens et al., 2023; Ludwig, et al., 2021; Zhang et al., 2021; Zhang et al., 2022). Connectivity-mapping studies revealed that GIPR^AP^ neurons project to and inhibit the activity of neighbouring excitatory GFRAL and GLP-1R neurons locally within the AP (Zhang et al., 2021; Zhang et al., 2022). A wealth of studies have demonstrated that GIPR^AP^ neuron activation decreases conditioned taste avoidance induced by GDF15, LiCl, GLP-1, and PYY through these local inhibitory circuits (Borner et al., 2021; Borner et al., 2023; Samms et al., 2022; Zhang et al., 2022). In the context of food intake, we propose that the orexigenic influence of GIPR^AP^ neurons is mediated through their tonic inhibition of satiation-promoting post synaptic partners. To this end we show that inhibition of GIPR^AP^ cells in conjunction with exendin-4 treatment enhances the GLP-1R anorectic effect through promoting meal termination. Work by Gutgesell and colleagues revealing that GIPR antagonism mimics the transcriptional responses of GLP-1R agonism in the DVC (Gutgesell et al., 2025; Liu et al., 2025) further supports a model whereby GIPR^AP^ neurons provide inhibitory tone to regulate both the aversive and satiating function of neighbouring anorectic neurons. In direct support of this model, we show that selective loss of *Gipr* in AP neurons attenuates the ability of pharmacological GIP antagonism to suppress appetite. This represents an important advancement in rationalizing the beneficial effects of both GIPR agonism and antagonism (Adriaenssens, 2025).

Our region-specific loss of function studies suggest that endogenous gut-derived GIP plays a role in the recruitment and anorectic effect of *Gipr*-expressing cells in the NTS. These findings are in accordance with recent studies showing that GIPR^NTS^ neurons are required for small intestinal lipids and metformin-mediated reduction in food intake in high-fat fed rats (Kuah et al., 2025). These anorectic effects are proposed to be mediated by increased GIP secretion caused by lipid infusions (Kuah et al., 2025). Complementary studies have shown that chemogenic activation of K-cells, and consequent GIP release, reduced food intake and body weight in diet induced obese mice (Lewis et al., 2024). Together, these findings imply that GIPR neurons in the NTS are able to induce an anorectic phenotype when recruited and that circulating GIP may play a crucial role.

Collectively, our data suggest that the net behavioural and therapeutic outcome of any GIP-based strategy is likely to reflect the balance of engagement across these anatomically segregated and functionally opposed circuits, rather than a single uniform central GIPR action. The ratio of AP:NTS GIPR circuit activation will vary with pharmacokinetic properties and central penetrance of different GIPR agonists/antagonists. Therefore, characterising the relative degree to which these opposing circuits are engaged by current and future obesity pharmacotherapies will inform our understanding of their mechanisms of action.

### Peripheral Inputs to GIPR AP and NTS Cells Encode Distinct Interoceptive Modalities

A defining feature of the DVC is its role as the first central relay within the brain for vagal interoceptive information. We found that peripheral inputs to GIPR^AP^ and GIPR^NTS^ neurons map onto molecularly defined vagal afferent classes encoding distinct sensory modalities. GIPR^NTS^ neurons were preferentially innervated by *Oxtr*+ afferents—associated with intraganglionic laminar endings and gastrointestinal mechanosensation—whereas GIPR^AP^ neurons received greater input from *Gpr65*+ mucosal afferents associated with luminal chemosensation (Bai et al., 2019; Williams et al., 2016). The apparent segregation of VAN input to GIPR brainstem populations was mirrored *in vivo*: oxytocin strongly recruited GIPR^NTS^ neurons, whereas chow, a hypercaloric mixed meal and liquid sucrose preferentially activated GIPR^AP^ neurons.

These observations suggest that in the physiological context of a meal, mechanically and chemically derived gut signals are parsed into anatomically distinct and functionally segregated GIPR hindbrain circuits. Mechanosensory, distension-encoding input is an established driver of meal termination, and its integrity is required to prevent hyperphagia and excess weight gain in response to energy-dense diets (McDougle et al., 2021). We find that this input is routed to the anorectic GIPR^NTS^ population, consistent with a role for these neurons in promoting satiation and satiety. Chemosensory input relaying nutrient exposure is instead routed to GIPR^AP^ neurons, whose orexigenic, aversion-gating output may serve to sustain ingestion of nutritive loads while constraining inappropriate malaise.

Such modality-based segregation could allow the hindbrain to dynamically balance nutrient acquisition against satiation and aversion within a single ingestive episode. We therefore hypothesize that GIPR^AP^ and GIPR^NTS^ neurons are recruited during different temporal phases of the prandial cycle, such that their opposing outputs are sequential. Future studies should confirm the temporal order in which mechanical and chemical signals are integrated across a meal by GIPR brainstem neurons.

### GIPR Brainstem Neurocircuits Remodel in Response to Obesogenic Challenge

Strikingly, we found that Western diet led to a robust vagal denervation of GIPR^NTS^ neurons, preferentially affecting *Oxtr*+ VANs. By contrast, vagal inputs to GIPR^AP^ neurons, which are primarily chemosensory, did not undergo maladaptive remodelling. These findings imply that, in obesogenic environments, central representation of gastrointestinal mechanosensation to GIPR^NTS^ neurons could be impaired, potentially leading to over-consumption and chronic differences in food intake patterns. Obesity is increasingly understood not merely as a disorder of energy imbalance but as a state of maladaptive neurocircuit reorganisation across distributed networks controlling appetite, reward, and interoception (de Lartigue et al., 2026; Ferrario et al., 2024). Obesogenic diets remodel hypothalamic and cortical feeding circuits, desensitising AgRP neurons to dietary fat and reshaping synaptic and structural connectivity in the lateral hypothalamus and orbitofrontal cortex (Beutler et al., 2020; Linehan et al., 2020; Thompson et al., 2017), and degrade vagal afferent function, blunting gastric-distension signalling and satiety responsiveness (Kentish et al., 2012; Loper et al., 2021; McDougle et al., 2021). They also attenuate central incretin responsiveness, altering the temporal profile and magnitude of GLP-1R–mediated hypophagia and diminishing sensitivity to GLP-1R agonists (Bales et al., 2022; Duca et al., 2013; Mul et al., 2013). Our data extend these principles to brainstem GIPR circuits and highlight that different vagal populations present varied susceptibility to this maladaptive rewiring.

## Limitations

Several limitations qualify our conclusions. Our circuit maps rely on rabies-based monosynaptic input tracing, which, even with the enhanced transfer and neuronal viability afforded by CVS-N2c(ΔG) systems (Reardon et al., 2016), provides a relative rather than absolute readout of connectivity. The WD-associated changes in input fraction we describe are therefore best interpreted as shifts in relative connectivity, rather than a direct readout of synaptic strength. In addition, our incomplete (∼50%) region-specific *Gipr* knockout likely underestimates the contribution of *Gipr* expression in AP/NTS neurons, and endogenous GIP signalling. A key finding from our study is that selective loss of *Gipr* expression in AP neurons attenuates GIPR antagonist-mediated appetite suppression. At the time of submission, a manuscript from co-authors on this study find hypothalamic GIPR-expressing neurons mediate the ability of GIPR antagonism to sensitize to the weight loss effects of GLP-1 receptor agonism. We rationalize these findings by acknowledging that GIPR^AP^ neurons may represent one of multiple GIPR circuits that contribute to the beneficial effects of GIPR antagonism. Finally, fibre photometry reports population-level dynamics and cannot resolve functional heterogeneity within AP or NTS GIPR neurons, which single-cell imaging may yet reveal.

## AKNOWLEDGMENTS

This study was supported by EFSD/Novo Nordisk Foundation Trust, grant awarded to A.E.A. N.F.B was additionally funded by the British Society for Neuroendocrinology, UCL RES scholarship and UCL Bogue Fellowship. Photometry experiments were supported by NIH grants R01-DK106399, R01-DK138127, and R01-DK145100 (Z.A.K.), and T32DK007007 and HHMI GT17734 (A.L.C). B.J. is supported by an MRC Clinician Scientist Fellowship (MR/Y00132X/1).

The authors would like to thank Tricia Tan for valuable discussions on GIPR pharmacology. The authors would like to thank Molly Strom and Petr Znamenskiy for kindly gifting us rabies helper virus.

## AUTHOR CONTRIBUTIONS

N.F.B. and A.E.A. conceived the project and designed research studies. N.F.B. led experiments. N.F.B., C.S., A.E.A. performed food intake experiments and data analysis. N.F.B., A.L.C. and K.C. performed photometry experiments and data analysis. N.F.B. and A.R. performed metabolic phenotyping experiments. N.F.B. conducted circuit tracing experiments. A.E.A.performed bioinformatic analysis. F.M.G., F.R., N.H., N.I., generated transgenic mice. I.D., B.J., D.I.B., S.T. provided research reagents. A.E.A. supervised the project.

N.F.B. and A.E.A. wrote the manuscript with input from all authors. For the purpose of open access, the author has applied a Creative Commons Attributions (CC BY) license to any Author Accepted Manuscript version arising from this submission.

## METHODS

### Key Resources Table

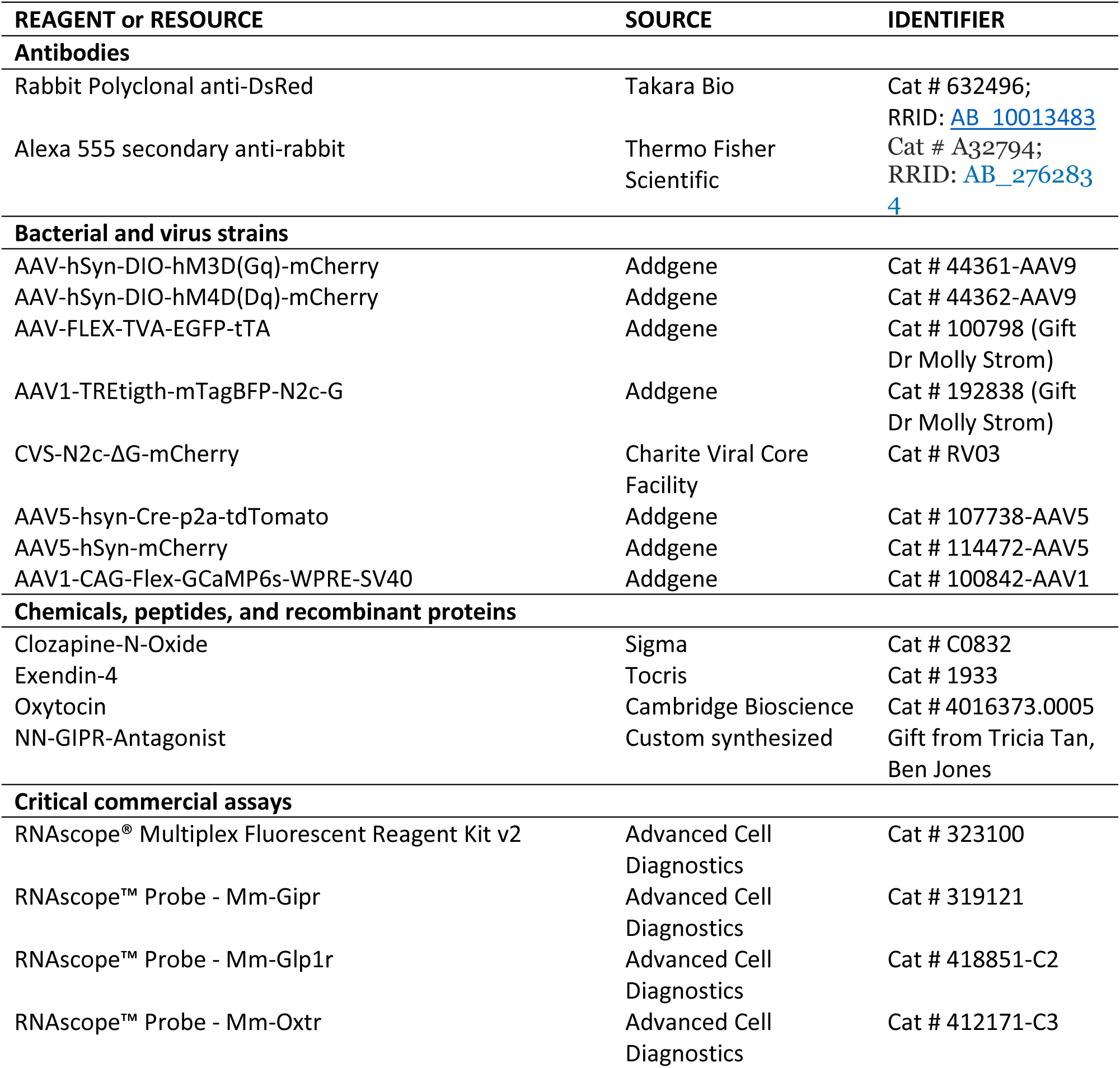
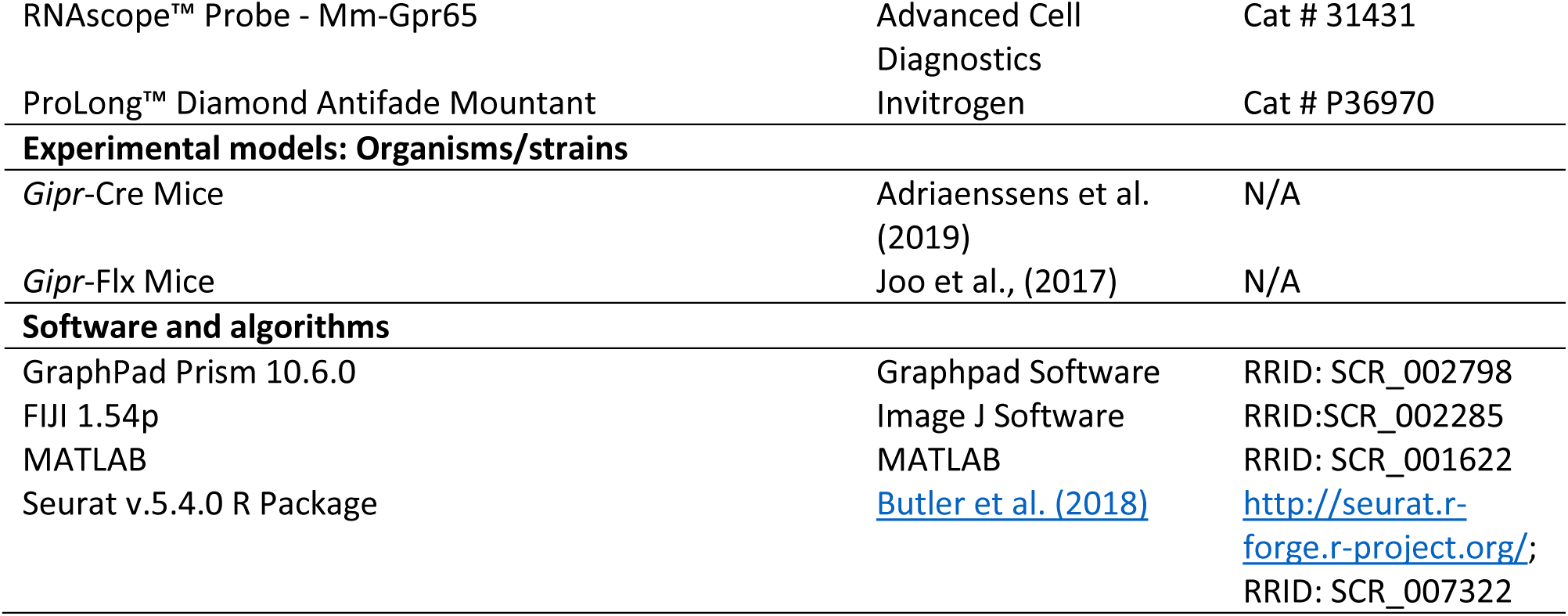

## Experimental Model and Subject Details

### Animals

All animal procedures conformed to the Animal (Scientific Procedures) Act 1986. Experimental protocols were approved by the UCL Animal Welfare and Ethical Review Body (Bloomsbury Campus). The studies were performed under the UK Home Office Project Licence, PP6526002. All mice were group-housed in individually ventilated cages with standard bedding and enrichment. Mice were housed in a temperature and humidity controlled room on a 12 h light/dark cycle (lights on 2:00, light out 14:00) with ad libitum access to water and standard chow diet unless stated otherwise. For studies performed in UCSF, experimental protocols were approved by the Institutional Animal Care and Use Committee of the University of California, San Francisco, following the National Institutes of Health guidelines for the Care and Use of Laboratory Animals.

### Mouse strains

For a subset of experiments *Gipr-*Cre mice (Adriaenssens et al., 2019) *ROSA*26-CAMPER (*Gipr-*Cre^CAMPER^) (Muntean et al., 2018) reporter strain to facilitate fluorescent detection of *Gipr-*expressing cells. Studies involving circuit tracing and *in vivo* phenotyping used animals that were heterozygous for both *Gipr* and Cre, and were label-free. For AP-and NTS-selective *Gipr* knock out studies, we used Giprfl/fl mice (Joo et al., 2017). All mouse lines were maintained on a mixed C57B6J/N genetic background.

## Method Details

### Stereotaxic Surgery

#### Brainstem viral injections

Animals were anaesthetized with ketamine hydrochloride (100 mg/kg) + medetomidine (10 mg/kg) and given meloxicam analgesia (5mg/kg). Depth of anaesthesia was determined by absence of pedal reflex and temperature was maintained using a homeothermic monitoring system. Animal was placed in a stereotaxic frame and ophthalmic ointment was applied to eyes. Head was ventroflexed such as the nose and neck were at a right angle. The scalp was incised from the occipital crest to the first vertebrae. Muscle layers were vivisected and retracted to expose atlanto-occipital membrane. Viral vectors were administered via pulled glass micropipettes into the NTS (0.2 mm AP, +-0.25 mm ML and-0.2 mm DV) or AP (0.65 mm AP, 0 mm ML and-0.16 mm DV) from obex. All viruses encoding chemogenetic effectors, Cre recombinase, or control reporters were administered in volumes of 75nl in NTS (bilateral injections) and 50nl in AP. Mice were allowed to recover for a minimum of 2 weeks before behavioural experiments. For rabies-assisted monosynaptic input tracing, 100 nl of a 1:1: mix of AAV-FLEX-TVA-EGFP-tTA and AAV1-TREtigth-mTagBFP-N2c-G were injected into the NTS or AP of Gipr-Cre mice. Fourteen days later, 100nl of CVS-N2c-ΔG-mCherry was administer in all injection sites. Mice were transcardially perfused for histology processing 4 days later.

#### Fiber photometry in the AP and NTS

Gipr-Cre mice were administered with AAV1-CAG-Flex-GCaMP6s via direct brainstem injections, as described above, into the AP or NTS. Mice were allowed to recover for 2 weeks before optic fiber implantation. Animals were anaesthetized with 2% isoflurane and place in a stereotaxic frame on a heating pad. Ophthalmic ointment was applied to the eyes and meloxicam was administered via subcutaneous injection (5 mg/kg) before surgery. Depth of surgical anaesthesia was determined by absence of pedal reflex. Local anaesthesia was applied (bupivacaine 0.25%) to the scalp and an incision was made through the midline to expose bregma, lambda sutures and occipital crest. A craniotomy was made using a dental drill (0.5mm). Optic fibre (Doric Lenses, MFC_400/430-0.48_6.5mm_MF2.5_FLT) and sleeve (Doric Lenses, SLEEVE_BR_2.5) were implanted at a 20 degree angle in the AP direction above the AP (1.45mm AP, 0 mm ML,-4.05 mm DV) or NTS (1.4mm AP, 0.3 mm ML,-4.15-4.2 mm DV) from the occipital crest. Mice were allowed to recover a minimum of 1 week before photometry experiments.

#### Viral vectors

AAVs were obtained from Addgene (AAV-hSyn-DIO-hM3D(Gq)-mCherry, 44361; AAV-hSyn-DIO-hM4D(Gi)-mCherry, 44362; AAV-CAG-Flex-GCaMP6s, 100842; AAV5-hsyn-Cre-p2a-tdTomato, 107738; AAV5-hSyn-mCherry, 114472), collaborators from the Crick Institute (AAV-FLEX-TVA-EGFP-tTA, AAV1-TREtigth-mTagBFP-N2c-G) and Berlin Viral Vector Core (CVS-N2c-ΔG-mCherry). Titers used were 2.6 x 10^12^ gc/ml for AAV-hSyn-DIO-hM3D(Gq)-mCherry and AAV-hSyn-DIO-hM4D(Gi)-mCherry, 1.5 x 10^12^ gc/ml for AAV5-hsyn-Cre-p2a-tdTomato and AAV5-hSyn-mCherry, 8.5 x 10^10^ gc/ml for AAV-FLEX-TVA-EGFP-tTA, 1.6 x 10^12^ gc/ml for AAV1-TREtigth-mTagBFP-N2c-G and 2.86 x 10^9^ gc/ml for CVS-N2c-ΔG-mCherry.

### Behavioural Studies

#### Food intake measurements

Food intake was monitored using open source Feeding experimental device 3 (FED3) pellet dispensers (Matikainen-Ankney et al., 2021) and 20mg Precision Pellets (TestDiet 5TUL/1811142). For intake measurements of the ad libitum eating paradigm, mice were fasted for 3 h in the light phase prior to lights off. Subsequently, chemogenetic mice were intraperitoneally (i.p.) injected with Clozapine N-oxide (CNO; Sigma) at a 1 mg/kg dose or saline vehicle 15 min prior to dark onset. For antagonist study, KO mice were injected subcutaneously (s.c.) with acyl-GIPR-antagonist (3µmol/kg) or saline vehicle 15 minutes prior to dark onset, FEDs were return at onset of dark phase. For post satiation paradigm, mice were fasted for 6hrs, which included the 1^st^ h of dark phase. They were then given ad lib access to food using FED3 devices and then injected i.p with CNO or saline vehicle. Food intake was monitored for the subsequent 24hrs. Knockout mice were not injected for baseline food intake studies. For light phase studies, mice were injected i.p. with CNO or saline vehicle 7 h after light on, changes were then monitored for 24hr. All chemogenetic experiments were conducted in DREADD expressing mice using a within-subject design. Food was returned at dark onset and monitored for a period of 24 h. For all food intake measurements, mice were habituated for 7 days to the FEDs and dosing before the start of testing. Meal pattern analysis was conducted from pellet data. A meal was defined empirically, using the biphasic distribution of inter-pellet intervals Jiang *et al,* 2025, as a sum of all bouts >0.04g with an intrameal interval of <1 min.

### Metabolic Phenotyping

#### EchoMRI

Gipr-Cre mice were exposed to ad libitum access western diet (Research Diets, D12079Bi) or standard rodent chow for 8-10 weeks. Mice received a body composition scan (EchoMRI™-100H) 4 weeks and 9 weeks after diet exposure to determine lean and fat mass.

#### Intraperitoneal glucose tolerance test (IPGTT)

Mice were fasted for 5h with ad libitum access to water prior to IPGTT. Mice received 1 g/kg glucose (20% glucose in sterile saline, i.p). Blood samples were collected from the tail vein before glucose injection for baseline measurement and then 15, 30, 60, 60 and 120 min after injection. Blood glucose levels were measured using a Roche ACCU-CHECK (Aviva) test kit. IPGTT tests were conducted 9 weeks after western diet exposure.

### Fiber Photometry

#### Behaviour

All fiber photometry recordings were performed in sound-isolated behaviour chambers (Coulbourn, Habitest Modular System; Med Associates, Davis Rig). Mice were habituated for overnight to the chambers prior to experiments. Handling and tethering habituation was also performed. Prior to each recording, photometry implants were cleaned with 70% ethanol using connector leaning sticks (MCC-S25). For all experiments, a baseline of 20 minutes was recorded before presentation or administration of a stimulus.

#### Photometry setup

Mice were tethered to a patch cable (Doric Lenses, MFP_400/460/900-0.48_2m_FCM-MF2.5). Continuous 6 mW blue LED (470 nm) and UV LED (405 nm) served as excitation light sources. These LEDs were driven by a multichannel hub (Thorlabs), modulated at 305 Hz and 505 Hz, respectively, and delivered to a filtered minicube (Doric Lenses, FMC6_AE(400-410)_E1(450-490)_F1(500-540)_E2(550-580)_ F2(600-680)_S) before connecting through optic fibres (Doric Lenses, MFP_400/460/900-0.48_2m_FCM-MF2.5). GCaMP calcium GFP signals and UV isosbestic signals were collected through the same fibres back to the dichroic ports of the minicube into a femtowatt silicon photoreceiver (Newport, 2151). Digital signals sampled at 1.0173 kHz were then demodulated, lock-in amplified and collected through a processor (RZ5P, Tucker-Davis Technologies). Data were then collected using the software Synapse (TDT), exported using Browser (TDT) and downsampled to 4 Hz in MATLAB before analysis.

GCaMP6s calcium signals acquired at 470 nm were normalized to the isosbestic control signal acquired at 405/415 nm using a linear regression fit derived from the baseline period, yielding a normalized fluorescence trace (Fnormalized). This normalized signal was then converted to a z-score using the mean and standard deviation of the baseline period preceding each stimulus: z = (Fnormalized − μ) / σ. For display purposes, mean traces were downsampled by a factor of 120 to reduce file size.

For most experiments, baseline activity was calculated from the 10 minutes preceding stimulus presentation, during which animals remained undisturbed in the behaviour chamber. The mean z-score during this baseline window was compared to the average z-score during the post-stimulus epoch of interest. Mean z-scores reported are for the entire post stimulus epoch, unless otherwise noted.

#### Histology

For immunohistochemistry and fluorescent in situ hybridization, animals were terminally anaesthetised with sodium pentobarbital (200 mg/kg) and transcardially perfused with heparinized 0.1M phosphate buffer saline (PBS) followed by 4% paraformaldehyde (PFA). Brains and nodose were extracted and post-fixed in 4% PFA at 4°C for 24 h. Tissue was then transferred to 30% sucrose solution for cryoprotection for 24 h.

#### Immunofluorescence labelling

Brains were coronally sectioned at a thickness of 30 μm using a cryostat and stored in cryoprotectant medium at-20°C. For immunofluorescent labelling, slices were washed in 0.1M PBS and then transferred to sodium citrate(10 mM, pH 6.0) for antigen retrieval at 80°C for 20 min. Tissue was washed in 0.1M PBS, blocked for 1h in 5% normal serum and then incubated in primary antibody (DsRed: Takara Bio, Cat number 632496, 1:1000; in blocking solution overnight at room temperature. Slices were washed and incubated in secondary antibody for 2 h at room temperature. Slices were washed and mounted on superfrost slides and coverslipped using Prolong diamond mounting medium (Invitrogen, Cat # P36970).

#### RNAscope in situ hybridization Nodose ganglia and brain slices

Nodose ganglia were sliced at 10 µm on a cryostat and collected on Supefrost Plus slides, then allowed to air dry at room temperature. Brain tissue was sliced at 16µm and collected on Superfrost slides. RNAscope was performed on sections using the RNAscope Multiplex Fluorescent Kit V.2 (Advanced Cell Diagnostics, Cat number: 323100) as per manufacturer’s instructions with a modification of the pretreatment procedure (Protease III incubation conducted for 20 min at 40°C) providing optimal signal detection of the target mRNAs. Probes for *Gpr65* (ACD Bio Cat number: 431431) *Oxtr* (ACD Bio Cat number: 412171), *Glp1r* (ACD Bio, Cat number: 418851) and *Gipr* (ACD Bio, Cat number: 319121) were hybridized and labelled with TSA vivid fluorophores. This was followed by additional processing for immunofluorescent labelling of viral reporter as described above. After completion, slides were cover slipped using Prolong Diamond mounting medium (Invitrogen, Cat # P36970).

#### Imaging

Imaging was performed on an EVOS M7000 multichannel epifluorescence imaging system (Invitrogen). Multichannel z-stack images were captured at 20x. Images were tiled and used to generate multichannel maximum intensity projection x-stacks using evos software. Images were processed using Image J analysis software.

### Transcriptomic Analysis

#### Mouse

Publicly-available transcriptomic data of single cells/nuclei from the mouse dorsal vagal complex produced by Zhang et al (Zhang et al., 2021a), Ludwig et al (Ludwig et al., 2021). *Gipr* positive neuronal clusters were identified and subsetted. ScDblFinder was used to identify doublets (Germain et al., 2021). The resulting datasets were re-normalized individually prior to merging and integration using Seurat (v.5.4.0) in R (Butler et al., 2018; Hao et al., 2021) as previously described (Huang et al., 2024). The integrated dataset was scaled and uniform manifold approximation and projection (UMAP) was generated. Neuronal clusters were assigned to the AP or NTS based on the expression patterns of signature genes previously identified (Ludwig et al., 2021) and confirmed with RNA in situ hybridization data from the Allen Brain Atlas. Co-expression of select targets in *Gipr* neurons was analysed and grouped by region ID using the Seurat DotPlot function.

Publicly-available transcriptomic data of single nuclei isolated from mouse nodose ganglia (Bai et al., 2019) was used to compare *Oxtr*+ versus *Gpr65*+ VAN populations.

#### Module score analysis of neurotransmitter identity

To characterise the neurotransmitter phenotype of neurons in the area postrema (AP) and nucleus tractus solitarius (NTS), per-cell module scores were computed for excitatory (glutamatergic), inhibitory (GABAergic) and glycinergic gene programmes using the AddModuleScore function in Seurat (v5) (Tirosh et al., 2016). Analyses were performed on the integrated single-nucleus RNA-sequencing object, restricted to neurons annotated as AP or NTS.

Gene sets were defined a priori from canonical neurotransmitter-identity markers and intersected with the genes detected in the dataset; only detected genes were retained for scoring. The gene sets were: excitatory/glutamatergic — *Slc17a6*, *Slc17a7*, *Slc17a8*; inhibitory/GABAergic — *Gad1*, *Gad2*, *Slc32a1*, *Reln, Lamp5, Npy, Tfap2a*; and glycinergic — *Slc6a5*, *Glra1*, *Glra2, Glra3, Glrb*. For each gene set, AddModuleScore computed the mean expression of the set in each cell minus the mean expression of a control gene set of equivalent size sampled from expression-level-matched bins (control parameter, ctrl = 25; random seed = 42).

Module scores were compared between regions and visualised as violin plots. Differences between AP and NTS were assessed using two-sided Wilcoxon rank-sum tests, with p-values adjusted for multiple comparisons across modules using the Benjamini–Hochberg method.

Data were visualised using ggplot2.

### Quantification and Statistical Analysis Data Analysis

All data are expressed as mean and SEM. Statistical analysis, detailed in figure legends, was performed using GraphPad Prism 10.6. Multiple comparisons were made using a 2-way-ANOVA with Sidak’s post hoc test. Single comparisons were made using a paired Student’s t-test. The number of mice or biological replicates used in each study is represented by the n number.

---

**Supplementary Figure 1:**
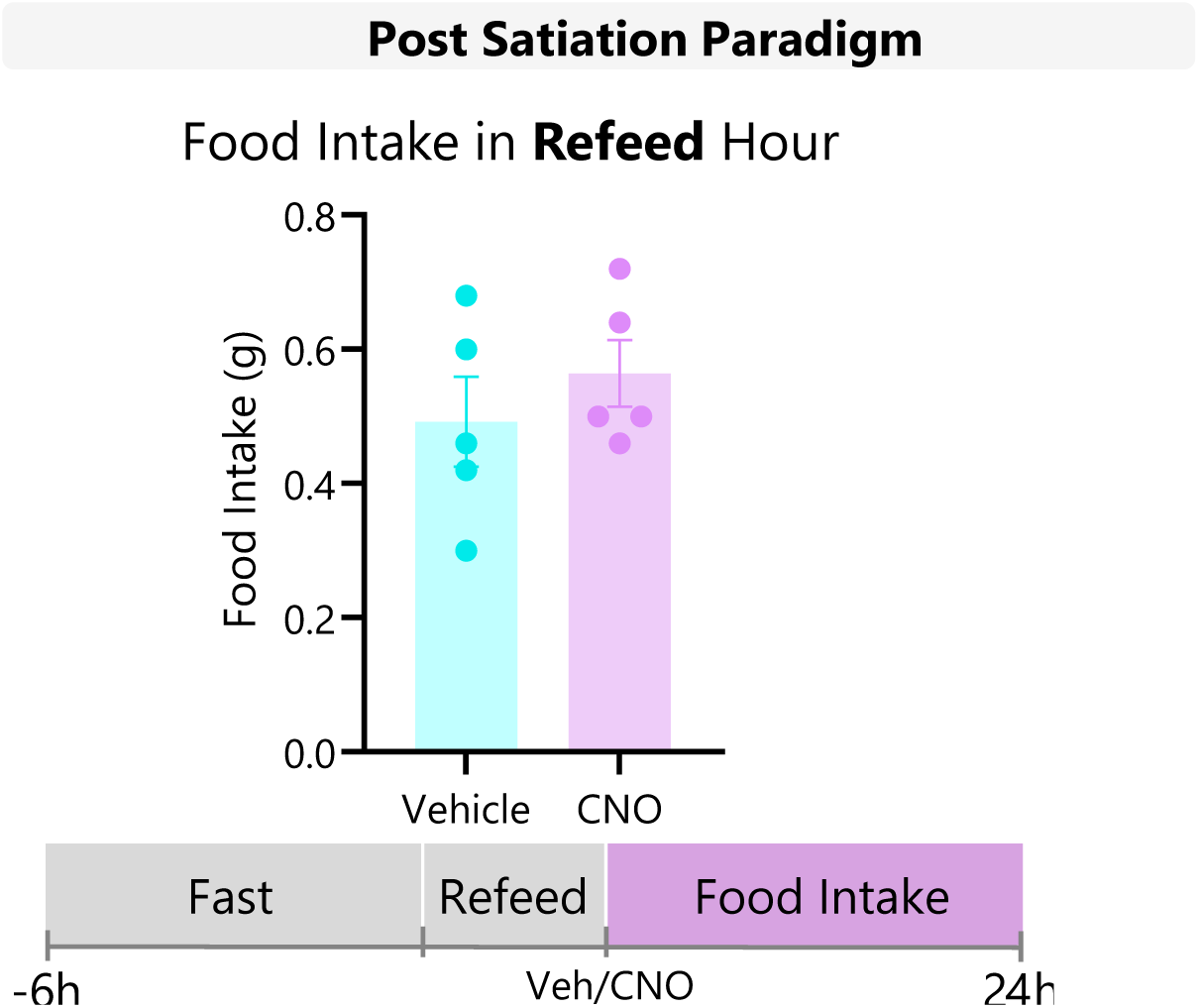
Post Satiation Paradigm: Mice were fasted for 6hr into the dark phase and given access to ad libitum feeding for an hour using FED3 devices. Food intake during refeeding period was measured to ensure no differences between treatment days. Clozapine-N-Oxide or Vehicle were intraperitoneally administer following the refeeding hour and food intake changes were then assessed in a sated background (n=5). Data are plotted as means +/-SEM. Statistical comparisons made using paired t-test as appropriate. * p < 0.05, ** p < 0.01, **** p < 0.001

**Supplementary Figure 2:**
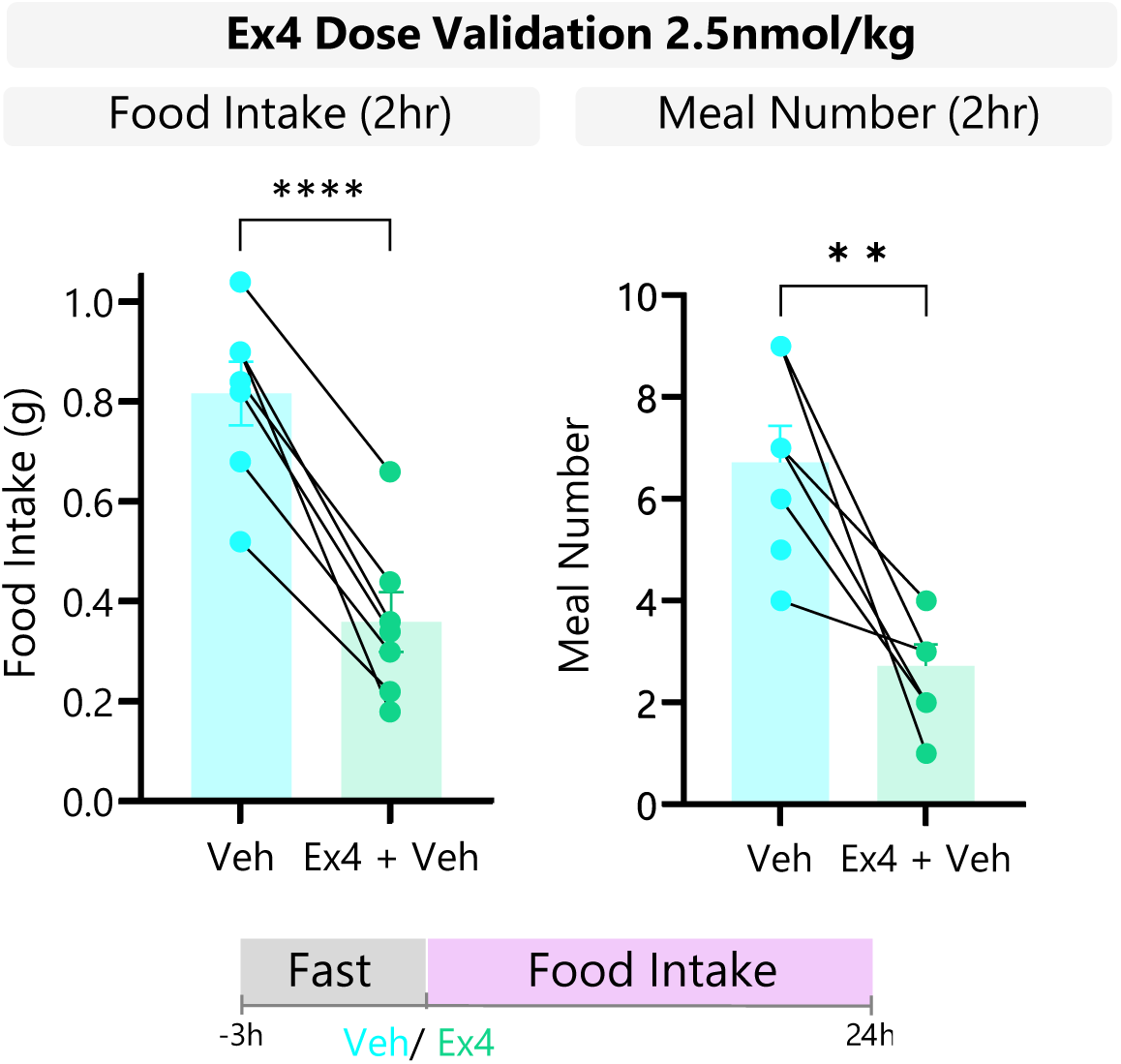
2.5 nmol/kg Exendin-4 elicits robust anorexia. Mice were dosed with vehicle of 2.5 nmol/Kg Exendin-4 (Ex-4) at the onset of the dark phase. Food intake was measured and quantified. Data are plotted as means +/-SEM. Statistical comparisons made using paired t-test as appropriate. * p < 0.05, ** p < 0.01, **** p < 0.001

**Supplementary Figure 3:**
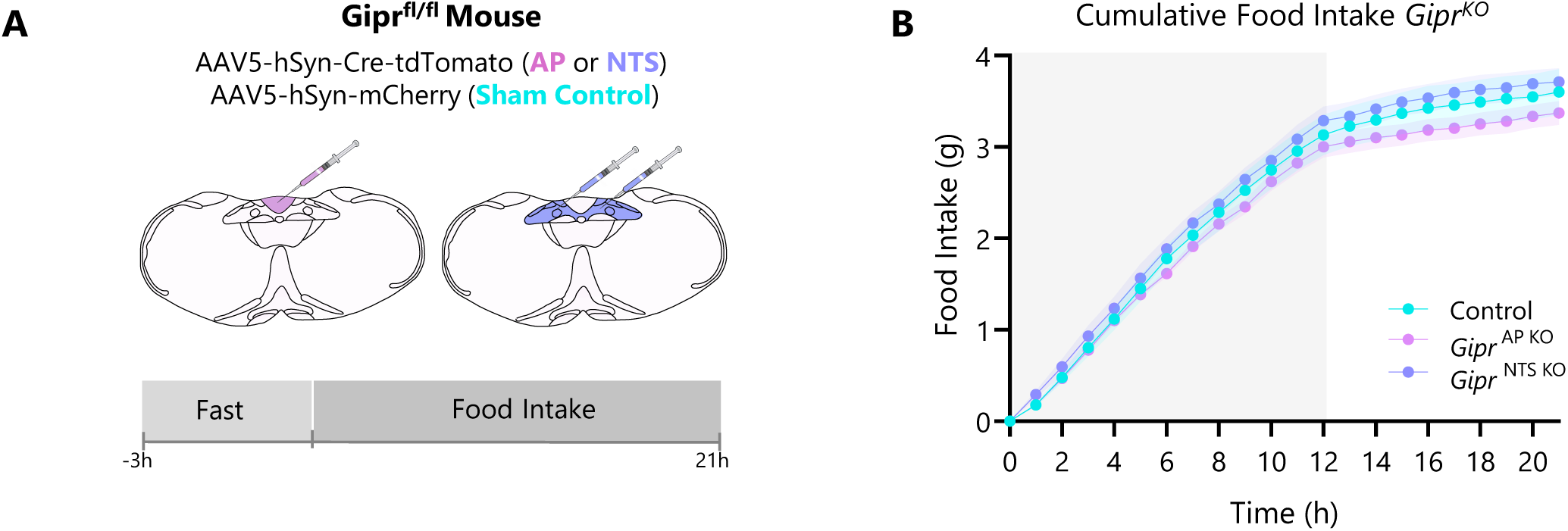
Loss of *Gipr* Expression in the AP and NTS Alters Feeding Behaviour. A-B, Gipr^fl/fl^ mice were injected with AAV-hSyn-Cre-tdTomato to produce selective KO of *Gipr* in the AP (Gipr^AP-KO^) or NTS (Gipr^NTS-KO^). Sham injected mice received a control virus in the AP and NTS. Baseline changes in food intake in dark phase feeding (B) were measured using FED3 devices in home cages. Mice were fasted for 3h of the light phase and FED3 devices were returned at onset of dark phase. Changes in food intake were assessed for 21hr. Data are plotted as means +/-SEM. Statistical comparisons made using a repeated measures 2-way ANOVA with a Sidak’s post-hoc test, paired or unpaired t-test as appropriate. * p < 0.05, ** p < 0.01, **** p < 0.001; n = 4-7

**Supplementary Figure 4:**
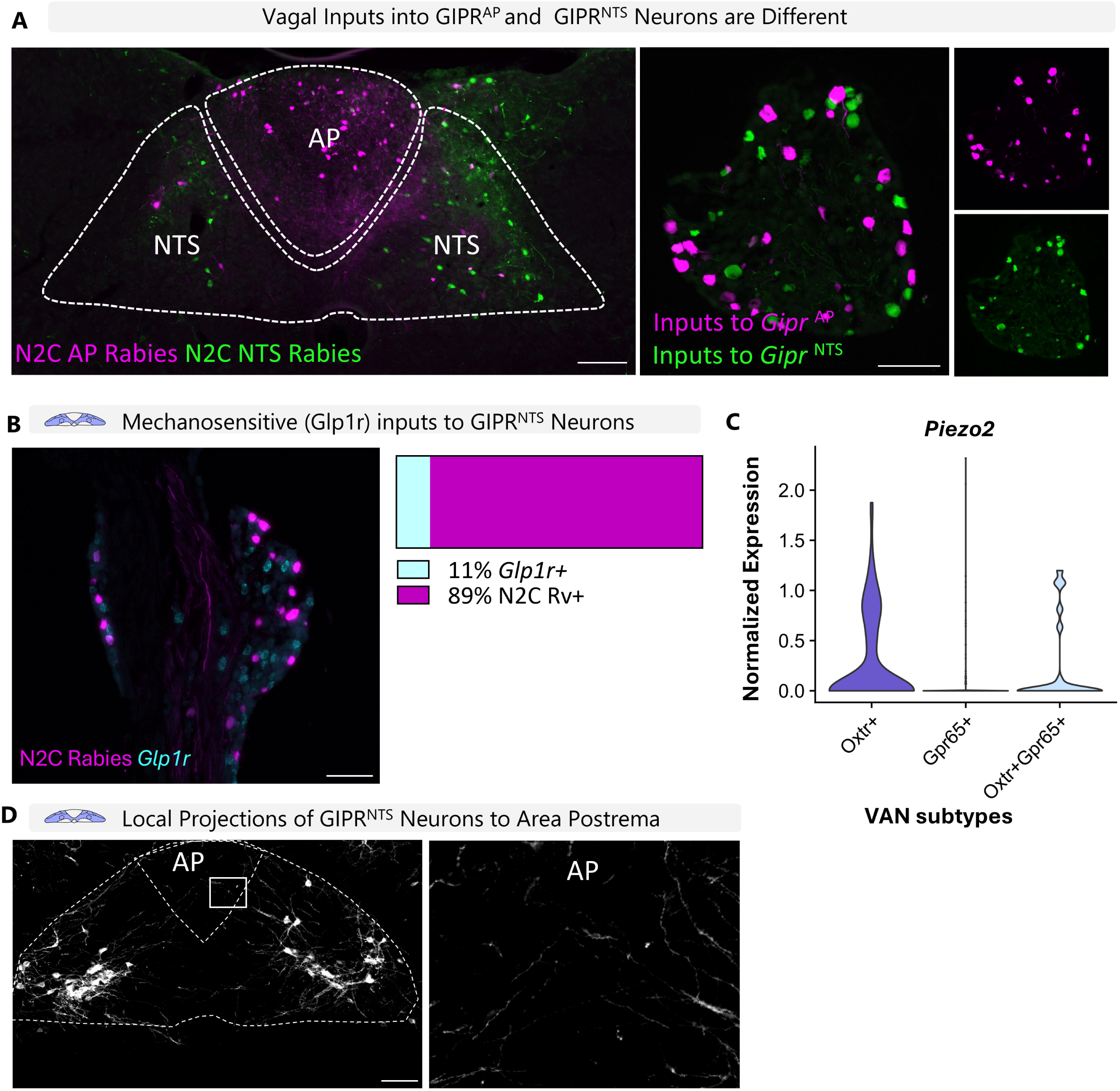
GIPR ^AP^ vs GIPR ^NTS^ Neurons Receive Distinct Peripheral Inputs and Exhibit Different Response Stimuli to Chemosensory Input. (A) Gipr-Cre mice were injected with AAV-TREtight-mTagBFP2-N2cG and AAV-hSyn-FLEX-splitTVA-EGFP-Tta helper virus into the AP and NTS. Subsequently, mice were injected with EnvA-N2C-mCherry rabies into the AP and EnVA-N2C-GFP rabies into the NTS. Peripheral inputs to AP and NTS neurons in same animals were largely non-overlapping. Image is representative from n=2 (B). (C) Publicly available snRNAseq data from murine nodose ganglia were analysed to quantify expression of the mechanosensor, *Piezo2*, across *Oxtr*+, *Gpr65*+, and *Oxtr*+/*Gpr65*+ vagal afferent neuron subtypes. (D) Gipr-Cre mice were injected with AAV-hM3Dq-mCherry into the NTS. Micrograph depicting processes of *Gipr* ^NTS^ neurons extending into the AP.

**Supplementary Figure 5:**
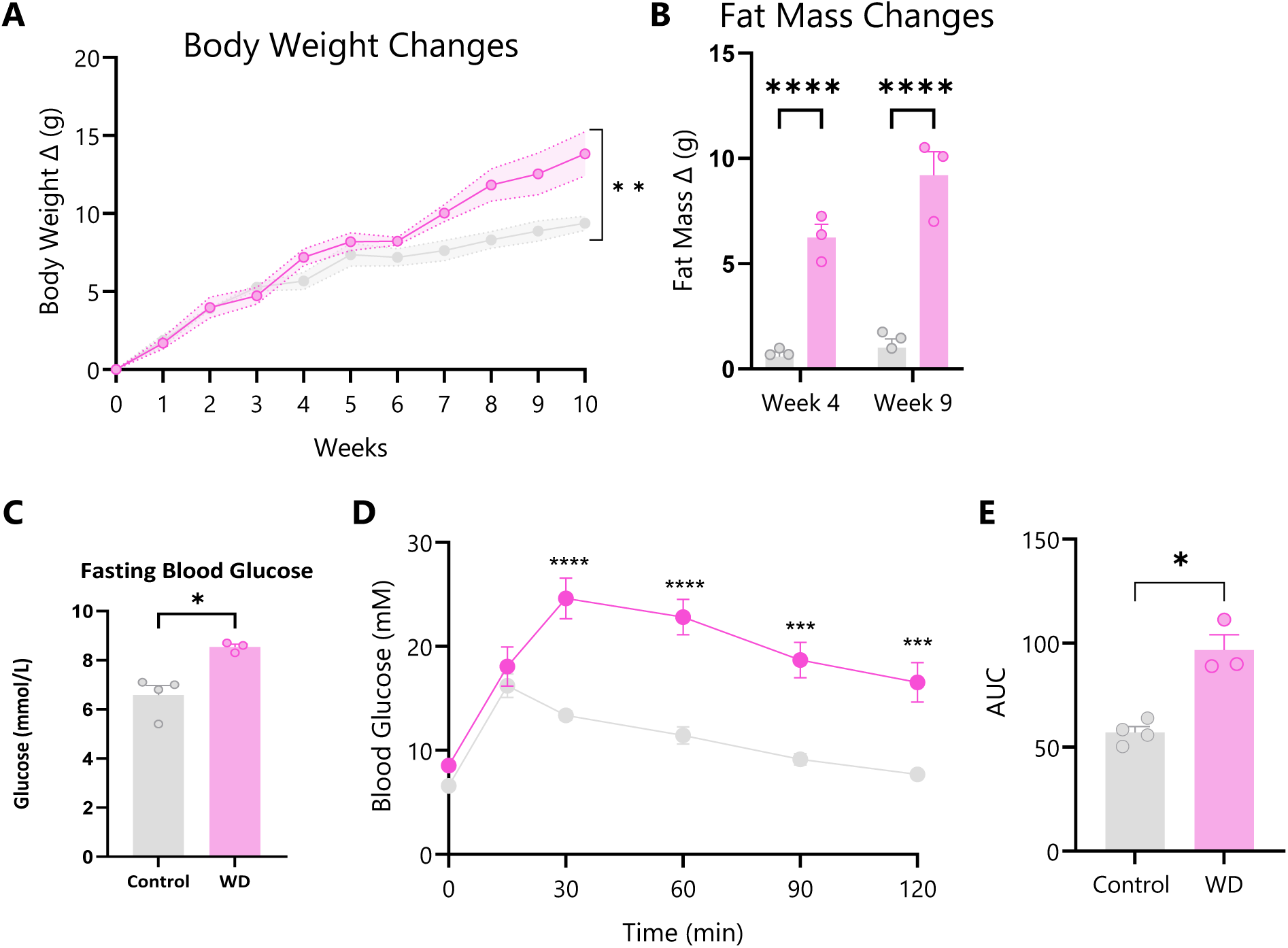
Metabolic Dysfunction Following Western Diet Exposure. Gipr-Cre mice were exposed to western diet or control diet for 8-10 weeks. (A) Body weight and (B) fat mass were measured and compared between control and western diet groups. (C-D) Changes in glucose tolerance was also measured. Data are plotted as means +/-SEM. Statistical comparisons made using a repeated measures 2-way ANOVA with a Sidak’s post-hoc test, paired or unpaired t-test as appropriate. * p < 0.05, ** p < 0.01, **** p < 0.001; n = 3-4

## REFERENCES

Adriaenssens, A., Biggs, E., Darwish, T., Tadross, J., Sukthankar, T., Girish, M.,…Reimann, F. (2019). Glucose-dependent insulinotropic polypeptide receptor-expressing cells in the hypothalamus regulate food intake. Cell Metabolism. 10.1016/j.cmet.2019.07.013

Adriaenssens, A., Broichhagen, J., de Bray, A., Ast, J., Hasib, A., Jones, B.,…Reimann, F. (2023). Hypothalamic and brainstem glucose-dependent insulinotropic polypeptide receptor neurons employ distinct mechanisms to affect feeding. JCI Insight, 8(10). 10.1172/jci.insight.164921

Adriaenssens, A. E. (2025). Unravelling the GIPR agonist versus antagonist paradox. Nature Metabolism, 7(6), 1111–1113. 10.1038/s42255-025-01299-6

Alexander, G. M., Rogan, S. C., Abbas, A. I., Armbruster, B. N., Pei, Y., Allen, J. A.,…Roth, B. L. (2009). Remote control of neuronal activity in transgenic mice expressing evolved G protein-coupled receptors. Neuron, 63(1), 27–39. 10.1016/j.neuron.2009.06.014

Bai, L., Mesgarzadeh, S., Ramesh, K. S., Huey, E. L., Liu, Y., Gray, L. A.,…Knight, Z. A. (2019). Genetic Identification of Vagal Sensory Neurons That Control Feeding. Cell, 179(5), 1129–1143.e1123. 10.1016/j.cell.2019.10.031

Bales, M. B., Centanni, S. W., Luchsinger, J. R., Fathi, P., Biddinger, J. E., Le, T. D. V.,…Ayala, J. E. (2022). High fat diet blunts stress-induced hypophagia and activation of Glp1r dorsal lateral septum neurons in male but not in female mice. Mol Metab, 64, 101571. 10.1016/j.molmet.2022.101571

Beutler, L. R., Corpuz, T. V., Ahn, J. S., Kosar, S., Song, W., Chen, Y., & Knight, Z. A. (2020). Obesity causes selective and long-lasting desensitization of AgRP neurons to dietary fat. Elife, 9. 10.7554/eLife.55909

Borner, T., Geisler, C. E., Fortin, S. M., Cosgrove, R., Alsina-Fernandez, J., Dogra, M.,…Hayes, M. R. (2021). GIP Receptor Agonism Attenuates GLP-1 Receptor Agonist-Induced Nausea and Emesis in Preclinical Models. Diabetes, 70(11), 2545–2553. 10.2337/db21-0459

Borner, T., Reiner, B. C., Crist, R. C., Furst, C. D., Doebley, S. A., Halas, J. G.,…Hayes, M. R. (2023). GIP receptor agonism blocks chemotherapy-induced nausea and vomiting. Mol Metab, 73, 101743. 10.1016/j.molmet.2023.101743

Browning, K. N., Verheijden, S., & Boeckxstaens, G. E. (2017). The Vagus Nerve in Appetite Regulation, Mood, and Intestinal Inflammation. Gastroenterology, 152(4), 730–744. 10.1053/j.gastro.2016.10.046

Butler, A., Hoffman, P., Smibert, P., Papalexi, E., & Satija, R. (2018). Integrating single-cell transcriptomic data across different conditions, technologies, and species. Nat Biotechnol, 36(5), 411–420. 10.1038/nbt.4096

Campbell, J. E., & Drucker, D. J. (2013). Pharmacology, physiology, and mechanisms of incretin hormone action. Cell Metab, 17(6), 819–837. 10.1016/j.cmet.2013.04.008

de Bray, A., Roberts, A. G., Armour, S., Tong, J., Huhn, C., Gatin-Fraudet, B.,…Hodson, D. J. (2025). Fluorescent GLP1R/GIPR dual agonist probes reveal cell targets in the pancreas and brain. Nature Metabolism, 7(8), 1536–1549. 10.1038/s42255-025-01342-6

de Lartigue, G., Brierley, D. I., & Choi, H. J. (2026). The critical role of gut-brain signalling in eating behaviour and obesity. Nat Rev Gastroenterol Hepatol. 10.1038/s41575-026-01203-x

Duca, F. A., Sakar, Y., & Covasa, M. (2013). Combination of obesity and high-fat feeding diminishes sensitivity to GLP-1R agonist exendin-4. Diabetes, 62(7), 2410–2415. 10.2337/db12-1204

Ellacott, K. L., Morton, G. J., Woods, S. C., Tso, P., & Schwartz, M. W. (2010). Assessment of feeding behavior in laboratory mice. Cell Metab, 12(1), 10–17. 10.1016/j.cmet.2010.06.001

Ferrario, C. R., Münzberg-Gruening, H., Rinaman, L., Betley, J. N., Borgland, S. L., Dus, M.,…Cooke, B. M. (2024). Obesity-and diet-induced plasticity in systems that control eating and energy balance. Obesity, 32(8), 1425–1440. 10.1002/oby.24060

Finan, B., Müller, T. D., Clemmensen, C., Perez-Tilve, D., DiMarchi, R. D., & Tschöp, M. H. (2016). Reappraisal of GIP Pharmacology for Metabolic Diseases. Trends Mol Med, 22(5), 359–376. 10.1016/j.molmed.2016.03.005

Frías, J. P., Davies, M. J., Rosenstock, J., Pérez Manghi, F. C., Fernández Landó, L., Bergman, B. K.,…Investigators, S.-. (2021). Tirzepatide versus Semaglutide Once Weekly in Patients with Type 2 Diabetes. N Engl J Med, 385(6), 503–515. 10.1056/NEJMoa2107519

Germain, P. L., Lun, A., Garcia Meixide, C., Macnair, W., & Robinson, M. D. (2021). Doublet identification in single-cell sequencing data using. F1000Res, 10, 979. 10.12688/f1000research.73600.2

Gutgesell, R. M., Khalil, A., Liskiewicz, A., Maity-Kumar, G., Novikoff, A., Grandl, G.,…Müller, T. D. (2025). GIPR agonism and antagonism decrease body weight and food intake via different mechanisms in male mice. Nature Metabolism, 7(6), 1282–1298. 10.1038/s42255-025-01294-x

Hao, Y., Hao, S., Andersen-Nissen, E., Mauck, W. M., Zheng, S., Butler, A.,…Satija, R. (2021). Integrated analysis of multimodal single-cell data. Cell, 184(13), 3573–3587.e3529. 10.1016/j.cell.2021.04.048

Holst, J. J., Orskov, C., Nielsen, O. V., & Schwartz, T. W. (1987). Truncated glucagon-like peptide I, an insulin-releasing hormone from the distal gut. FEBS Lett, 211(2), 169–174. 10.1016/0014-5793(87)81430-8

Huang, K. P., Acosta, A. A., Ghidewon, M. Y., McKnight, A. D., Almeida, M. S., Nyema, N. T.,…Alhadeff, A. L. (2024). Dissociable hindbrain GLP1R circuits for satiety and aversion. Nature, 632(8025), 585–593. 10.1038/s41586-024-07685-6

Jastreboff, A. M., Ryan, D. H., Bays, H. E., Ebeling, P. R., Mackowski, M. G., Philipose, N.,…Pannacciulli, N. (2025). Once-Monthly Maridebart Cafraglutide for the Treatment of Obesity - A Phase 2 Trial. N Engl J Med, 393(9), 843–857. 10.1056/NEJMoa2504214

Joo, E., Harada, N., Yamane, S., Fukushima, T., Taura, D., Iwasaki, K.,…Inagaki, N. (2017). Inhibition of Gastric Inhibitory Polypeptide Receptor Signaling in Adipose Tissue Reduces Insulin Resistance and Hepatic Steatosis in High-Fat Diet-Fed Mice. Diabetes, 66(4), 868–879. 10.2337/db16-0758

Jänig, W. (1996). Neurobiology of visceral afferent neurons: neuroanatomy, functions, organ regulations and sensations. Biol Psychol, 42(1-2), 29–51. 10.1016/0301-0511(95)05145-7

Kentish, S., Li, H., Philp, L. K., O’Donnell, T. A., Isaacs, N. J., Young, R. L.,…Page, A. J. (2012). Diet-induced adaptation of vagal afferent function. J Physiol, 590(1), 209–221. 10.1113/jphysiol.2011.222158

Kuah, R., Wang, M. T., Yang, Z., Back, G., Li, R. J. W., Bruce, K.,…Lam, T. K. T. (2025). Metformin Boosts Intestinal Lipid Sensing via GIP to Suppress Feeding. Diabetes, 74(11), 1906–1917. 10.2337/db25-0100

Lauritsen, K. B., Moody, A. J., Christensen, K. C., & Lindkaer Jensen, S. (1980). Gastric inhibitory polypeptide (GIP) and insulin release after small-bowel resection in man. Scand J Gastroenterol, 15(7), 833–840. 10.3109/00365528009181538

Lewis, J. E., Nuzzaci, D., James-Okoro, P. P., Montaner, M., O’Flaherty, E., Darwish, T.,…Reimann, F. (2024). Stimulating intestinal GIP release reduces food intake and body weight in mice. Mol Metab, 84, 101945. 10.1016/j.molmet.2024.101945

Linehan, V., Fang, L. Z., Parsons, M. P., & Hirasawa, M. (2020). High-fat diet induces time-dependent synaptic plasticity of the lateral hypothalamus. Mol Metab, 36, 100977. 10.1016/j.molmet.2020.100977

Liskiewicz, A., Khalil, A., Liskiewicz, D., Novikoff, A., Grandl, G., Maity-Kumar, G.,…Müller, T. D. (2023). Glucose-dependent insulinotropic polypeptide regulates body weight and food intake via GABAergic neurons in mice. Nat Metab, 5(12), 2075–2085. 10.1038/s42255-023-00931-7

Liu, C. M., Killion, E. A., Hammoud, R., Lu, S.-C., Komorowski, R., Liu, T.,…Véniant, M. M. (2025). GIPR-Ab/GLP-1 peptide–antibody conjugate requires brain GIPR and GLP-1R for additive weight loss in obese mice. Nature Metabolism, 7(6), 1266–1281. 10.1038/s42255-025-01295-w

Loper, H., Leinen, M., Bassoff, L., Sample, J., Romero-Ortega, M., Gustafson, K. J.,…Schiefer, M. A. (2021). Both high fat and high carbohydrate diets impair vagus nerve signaling of satiety. Sci Rep, 11(1), 10394. 10.1038/s41598-021-89465-0

Ludwig, M. Q., Cheng, W., Gordian, D., Lee, J., Paulsen, S. J., Hansen, S. N.,…Pers, T. H. (2021). A genetic map of the mouse dorsal vagal complex and its role in obesity. Nat Metab, 3(4), 530–545. 10.1038/s42255-021-00363-1

Ludwig, M. Q., Todorov, P. V., Egerod, K. L., Olson, D. P., & Pers, T. H. (2021). Single-Cell Mapping of GLP-1 and GIP Receptor Expression in the Dorsal Vagal Complex. Diabetes, 70(9), 1945–1955. 10.2337/dbi21-0003

Matikainen-Ankney, B. A., Earnest, T., Ali, M., Casey, E., Wang, J. G., Sutton, A. K.,…Kravitz, A. V. (2021). An open-source device for measuring food intake and operant behavior in rodent home-cages. Elife, 10. 10.7554/eLife.66173

McDougle, M., Quinn, D., Diepenbroek, C., Singh, A., de la Serre, C., & de Lartigue, G. (2021). Intact vagal gut-brain signalling prevents hyperphagia and excessive weight gain in response to high-fat high-sugar diet. Acta Physiologica, 231(3), e13530. 10.1111/apha.13530

Mul, J. D., Begg, D. P., Barrera, J. G., Li, B., Matter, E. K., D’Alessio, D. A.,…Sandoval, D. A. (2013). High-fat diet changes the temporal profile of GLP-1 receptor-mediated hypophagia in rats. Am J Physiol Regul Integr Comp Physiol, 305(1), R68–77. 10.1152/ajpregu.00588.2012

Muntean, B. S., Zucca, S., MacMullen, C. M., Dao, M. T., Johnston, C., Iwamoto, H.,…Martemyanov, K. A. (2018). Interrogating the Spatiotemporal Landscape of Neuromodulatory GPCR Signaling by Real-Time Imaging of cAMP in Intact Neurons and Circuits. Cell Rep, 24(4), 1081–1084. 10.1016/j.celrep.2018.07.031

Müller, T. D., Adriaenssens, A., Ahrén, B., Blüher, M., Birkenfeld, A. L., Campbell, J. E.,…Tschöp, M. H. (2025). Glucose-dependent insulinotropic polypeptide (GIP). Mol Metab, 95, 102118. 10.1016/j.molmet.2025.102118

Müller, T. D., Clemmensen, C., Finan, B., DiMarchi, R. D., & Tschöp, M. H. (2018). Anti-Obesity Therapy: from Rainbow Pills to Polyagonists. Pharmacol Rev, 70(4), 712–746. 10.1124/pr.117.014803

Page, A. J., & Kentish, S. J. (2017). Plasticity of gastrointestinal vagal afferent satiety signals. Neurogastroenterology & Motility, 29(5), e12973. 10.1111/nmo.12973

Pool, A. H., Poldsam, H., Chen, S., Thomson, M., & Oka, Y. (2023). Recovery of missing single-cell RNA-sequencing data with optimized transcriptomic references. Nat Methods, 20(10), 1506–1515. 10.1038/s41592-023-02003-w

Rathod, Y. D., & Di Fulvio, M. (2021). The feeding microstructure of male and female mice. PLoS One, 16(2), e0246569. 10.1371/journal.pone.0246569

Reiner, B. C., Crist, R. C., Borner, T., Doyle, R. P., Hayes, M. R., & De Jonghe, B. C. (2022). Single nuclei RNA sequencing of the rat AP and NTS following GDF15 treatment. Mol Metab, 56, 101422. 10.1016/j.molmet.2021.101422

Samms, R. J., Cosgrove, R., Snider, B. M., Furber, E. C., Droz, B. A., Briere, D. A.,…Ai, M. (2022). GIPR Agonism Inhibits PYY-Induced Nausea-Like Behavior. Diabetes. 10.2337/db21-0848

Thompson, J. L., Drysdale, M., Baimel, C., Kaur, M., MacGowan, T., Pitman, K. A., & Borgland, S. L. (2017). Obesity-Induced Structural and Neuronal Plasticity in the Lateral Orbitofrontal Cortex. Neuropsychopharmacology, 42(7), 1480–1490. 10.1038/npp.2016.284

Tirosh, I., Izar, B., Prakadan, S. M., Wadsworth, M. H., Treacy, D., Trombetta, J. J.,…Garraway, L. A. (2016). Dissecting the multicellular ecosystem of metastatic melanoma by single-cell RNA-seq. Science, 352(6282), 189–196. 10.1126/science.aad0501

Williams, Erika K., Chang, Rui B., Strochlic, David E., Umans, Benjamin D., Lowell, Bradford B., & Liberles, Stephen D. (2016). Sensory Neurons that Detect Stretch and Nutrients in the Digestive System. Cell, 166(1), 209–221. 10.1016/j.cell.2016.05.011

Yang, B., Gelfanov, V. M., El, K., Chen, A., Rohlfs, R., DuBois, B.,…Finan, B. (2022). Discovery of a potent GIPR peptide antagonist that is effective in rodent and human systems. Mol Metab, 66, 101638. 10.1016/j.molmet.2022.101638

Young, M. D., & Behjati, S. (2020). SoupX removes ambient RNA contamination from droplet-based single-cell RNA sequencing data. Gigascience, 9(12). 10.1093/gigascience/giaa151

Zhang, C., Kaye, J. A., Cai, Z., Wang, Y., Prescott, S. L., & Liberles, S. D. (2021b). Area Postrema Cell Types that Mediate Nausea-Associated Behaviors. Neuron, 109(3), 461–472.e465. 10.1016/j.neuron.2020.11.010

Zhang, C., Vincelette, L. K., Reimann, F., & Liberles, S. D. (2022). A brainstem circuit for nausea suppression. Cell Reports, 39(11). 10.1016/j.celrep.2022.110953

Zhang, Q., Delessa, C. T., Augustin, R., Bakhti, M., Colldén, G., Drucker, D. J.,…Müller, T. D. (2021). The glucose-dependent insulinotropic polypeptide (GIP) regulates body weight and food intake via CNS-GIPR signaling. Cell Metab. 10.1016/j.cmet.2021.01.015

Zorrilla, E. P., Inoue, K., Fekete, E. M., Tabarin, A., Valdez, G. R., & Koob, G. F. (2005). Measuring meals: structure of prandial food and water intake of rats. Am J Physiol Regul Integr Comp Physiol, 288(6), R1450–1467. 10.1152/ajpregu.00175.2004

